# The type III-B CRISPR-Cas System Affects Energy Metabolism and Adaptation in the Archaeon *Saccharolobus solfataricus*

**DOI:** 10.1101/2024.09.02.610847

**Authors:** Erika Wimmer, Logan H. Hodgskiss, Melina Kerou, Isabelle A. Zink, Christa Schleper

## Abstract

Type III CRISPR-Cas immune systems that recognize and cleave extrachromosomal RNA when active, are particularly widespread in archaea. Mechanistically, these systems have the potential to regulate gene expression of host genes on a post-transcriptional level, but very little is known about any potential accessory roles of type III-B systems beyond immunity. We have created knockout mutants of a type III-B CRISPR-Cas complex in the thermoacidophilic archaeon *Saccharolobus solfataricus* to investigate potential secondary functions of the type III-B system. Deletion mutants exhibited an accelerate growth but were less quickly adaptable to changes in carbon sources in their growth media. In line with this phenotype, upregulated genes were significantly enriched in functional categories of energy production and conversion, as well as with carbohydrate or amino acid transport and metabolism in RNAseq studies. Generally, a significant accumulation of genes encoding transmembrane proteins in the upregulated proportion of the transcriptome suggests interconnections between the type III-B CRISPR-Cas system and various membrane-associated processes. Notably, the deletion mutants did not lose their general virus- or plasmid defense activities indicating that this particular system might have been partially adopted for cellular regulatory roles.

## INTRODUCTION

CRISPR-Cas (Clustered Regularly Interspaced Short Palindromic Repeats - CRISPR associated proteins) immune systems are present in roughly a third of fully sequenced bacterial and the majority of archaeal genomes^1^. Despite the impressive diversity of known CRISPR-Cas systems, the principle first steps of the immune reaction remain: A small crRNA (CRISPR RNA) is transcribed from a so-called spacer in a CRISPR array in the host genome. Spacers are DNA snippets of previously encountered viruses and plasmids that have been stored in the host genome during the process of CRISPR adaptation^2,3^. A CRISPR-Cas protein effector forms a complex with one small crRNA to identify complementary nucleic acid targets (so-called protospacers) on invading genetic elements, such as viruses or plasmids^4^. Dependent on the CRISPR-Cas system, the bound nucleic acid target is then either directly cleaved by the Cas enzyme or abortive infection pathways are activated to clear the invader from the population^5^.

While most CRISPR-Cas effectors target double stranded DNA, type III and type VI CRISPR-Cas systems exhibit dedicated RNA-targeting mechanisms^6^. Type III CRISPR-Cas systems are frequently found in various archaeal phyla and are particularly abundant among Thermoproteota (formerly Crenarchaeota)^5^. These systems are especially intriguing because they are multi-subunit complexes that execute their immune reaction in three different ways: Upon recognition and binding of a specific RNA target (i.e. nascent mRNA transcribed from a plasmid), the Cas7 backbone of the type III complex cleaves the RNA protospacer. Furthermore, binding to the protospacer activates the Cas10 subunit of the complex which cleaves single stranded DNA in transcription bubbles promiscuously via its HD domain^7–10^. Concomitantly, the cyclase (palm) domain of Cas10 synthesizes cyclic molecules from ATP (cOAs)^11–14^ which in turn activate ancillary CARF-domain containing proteins which can be either RNases, DNases or proteases that indiscriminately degrade their substrates in the cell^15,16^. This leads to cell death or cell dormancy which curtails the spread of the invader in the population^17–23^.

Interestingly, the nucleolytic and abortive infection pathways of the type III CIRSPR-Cas complex can be inactivated, thereby only allowing specific cleavage of the RNA target. Complementarity of a sequence stretch at the 3’ end of the protospacer (the so called antitag) to the 5’ part of the crRNA or mismatched bases in certain parts of the crRNA:target duplex allosterically inhibit the catalytic activities of Cas10^8,9,11–14,24–26^. Therefore, type III CRISPR-Cas effector complexes can specifically cleave mRNA targets, which raises the question as to whether they also contribute to the regulation of host gene expression. Indeed, there are some indications for additional roles of type III CRISPR-Cas complexes apart from immunity. For instance, type III loci often contain high numbers of accessory genes potentially constituting a functional connection to cellular processes other than defense^27,28^. Also, in many cases type III CRISPR-*cas* operons are not genomically linked to CRISPR arrays or modules involved in the acquisition of new spacers, thus they are reliant on co-occurring CRISPR-Cas systems for a canonical immune response^5,29^. Furthermore, we find type III CRISPR-Cas systems to be among the most highly expressed genes in *Saccharolobus solfataricus*, also when no plasmid or virus invader is present.

*Sa. solfataricus* P1 belongs to the archaeal phylum Thermoproteota and contains two type III CRISPR-Cas complexes, a type III-D (Csm) and a type III-B (Cmr-ß), as well as three DNA-targeting type I-A complexes^5,30^. In vitro studies on the Cmr-ß complex from a close relative, *Sa. islandicus* (SisCmr-ß), revealed that catalytic activities of the complex are modulated by the additional subunit Cmr7^31,32^ as its presence coincided with starkly reduced cOA production by the Cas10 subunit^33^. In this study, the endogenous type I CRISPR-Cas system was harnessed to generate a *cmr-ß* knockout strain of *Sa. solfataricus* in order to discern its putative links to other processes and alternative roles besides immunity. A wide range of effects of the genomic deletion on energy metabolism and transport mechanisms indicate a general impact on the host’s physiology as revealed by RNAseq. In addition, knockout strains were more sensitive to switches from complex into single sugars media, indicating a role of the Cmr-ß complex in metabolic adaptability. Furthermore, we did not observe any effect on plasmid immunity upon type III CRISPR-Cas deletion, indicating that the Cmr-ß might not be a main contributor to type III CRISPR-Cas immune defense in *Sa. solfataricus*.

## RESULTS

### Type IA-CRISPR-Cas-mediated knockout of the type III-B CRISPR-*cas* gene module in *Sa. solfataricus* P1

The endogenous DNA-targeting type I-A CRISPR-Cas system was harnessed for deletion of the genes encoding the type III-B CRISPR-Cas complex Cmr-ß (SSOP1_RS10085 -SSOP1_RS10115) (Fig. 1A). Two different 37-bp protospacers with 3’ protospacer adjacent motif (PAM) 5’-NGG^34^ were selected within each of the three targeted genes (*cmr7, cmr6, cmr3/cas5*) on the *Sa. solfataricus* chromosome (Fig. 1A). In total, six different mini CRISPR arrays (miniCR) were assembled as described previously^35^ containing one spacer or a combination of two to three spacers targeting the respective genes on either coding or template strand (cs or ts, respectively). The miniCRs and two approximately 800-bp donor sequences for DNA repair corresponding to upstream and downstream flanking regions of the *cmr-ß* locus were cloned into the plasmid-based vector pIZ-GW^36^ (Fig. 1B). *Sa. solfataricus* M18 cells transformed with the genome-editing plasmids were enriched under selection in liquid medium. Although all constructs yielded the intended deletion as detected by culture PCR, one knockout culture was chosen for plating to isolate single colonies (Suppl. Fig. S1). The genomes of two purified knockout mutant cultures (KO1 and KO2) grown from two single colonies were sequenced showing that in comparison to the deposited reference (NZ_LT549890.1) the *cmr-ß* genes and their upstream basal promoter elements had been scarlessly excised from the genome in both mutants (Fig. 2). Interestingly, KO2 had additionally acquired a larger deletion (over 37 kb) between the genes SSOP1_RS14925 and SSOP1_RS15080, both of which are encoding IS5 family transposases (Suppl. tab. 1, Fig. 3).

**Figure 1.**
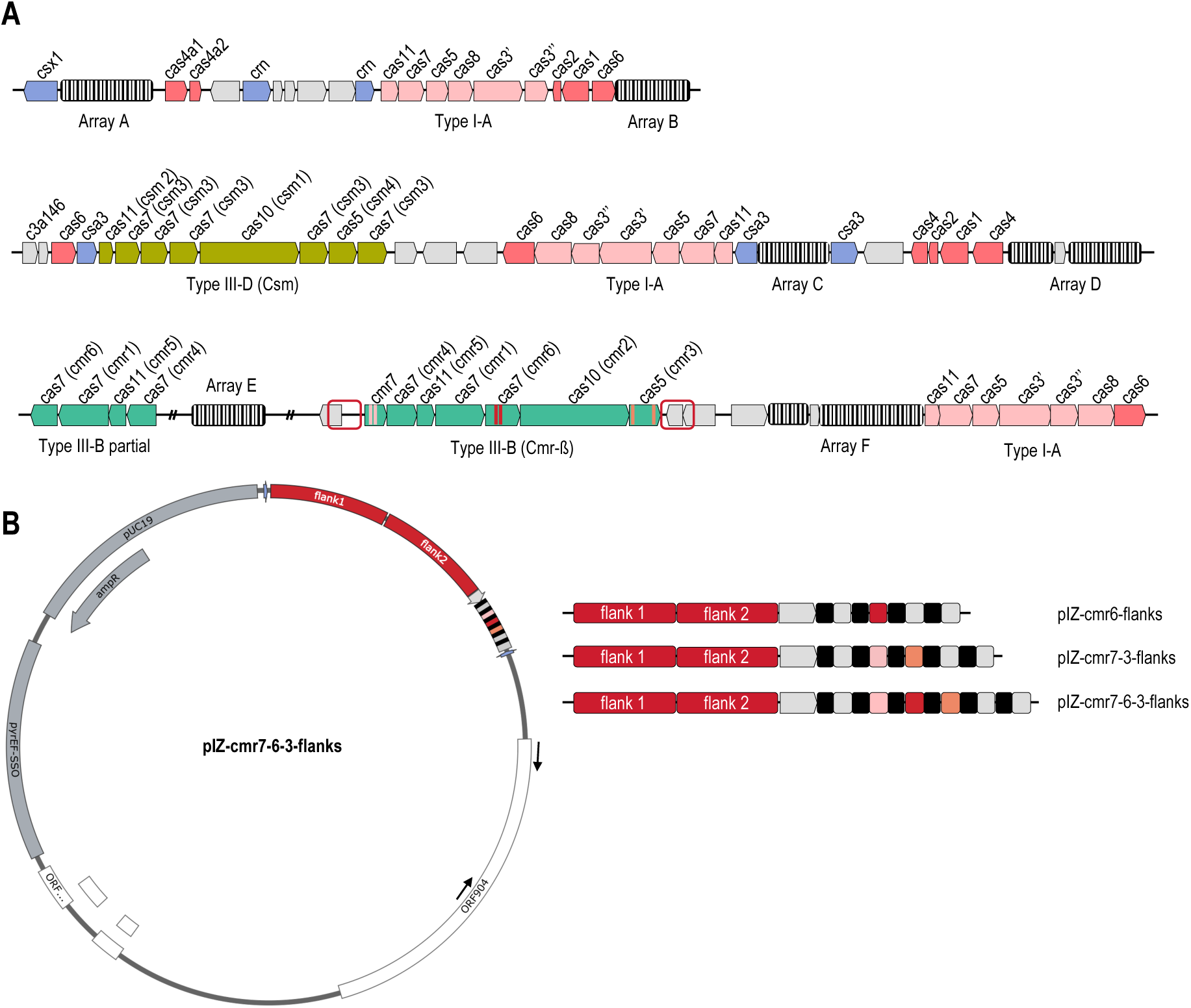
**(A)** Overview of the CRISPR-Cas systems in *Sa. solfataricus* P1. Three type I-A (pink), one type III-D (lime green), and one type III-B (green) system are present. In addition, a partial type III-B *cas* gene cluster can be detected. The strain harbors six CRISPR arrays (striped boxes), two are split through the insertion of fragments, in the case of array F stemming from a pNOB8-like conjugative plasmid. Six different protospacers that were chosen for the type I-A CRISPR-mediated deletion of the *cmr-ß* gene module are indicated. Two protospacers each are located in genes *cmr7* (pink), *cmr6* (red) and *cmr3* (salmon). Flanking sequences of the *cmr-ß* genes (red boxes) are provided as donor for the DNA repair process (see B). **(B)** Different combinations of artificial spacers (single to triple) all targeting either template or coding strand of the respective genes were cloned into a miniCR array on the plasmid-based vector pIZ, preceded by donor DNA sequences (red). Color code for miniCR: self-targeting spacers – equivalent to (A); non-targeting spacers (used for cloning procedure)– grey; repeats – black.

**Figure 2.**
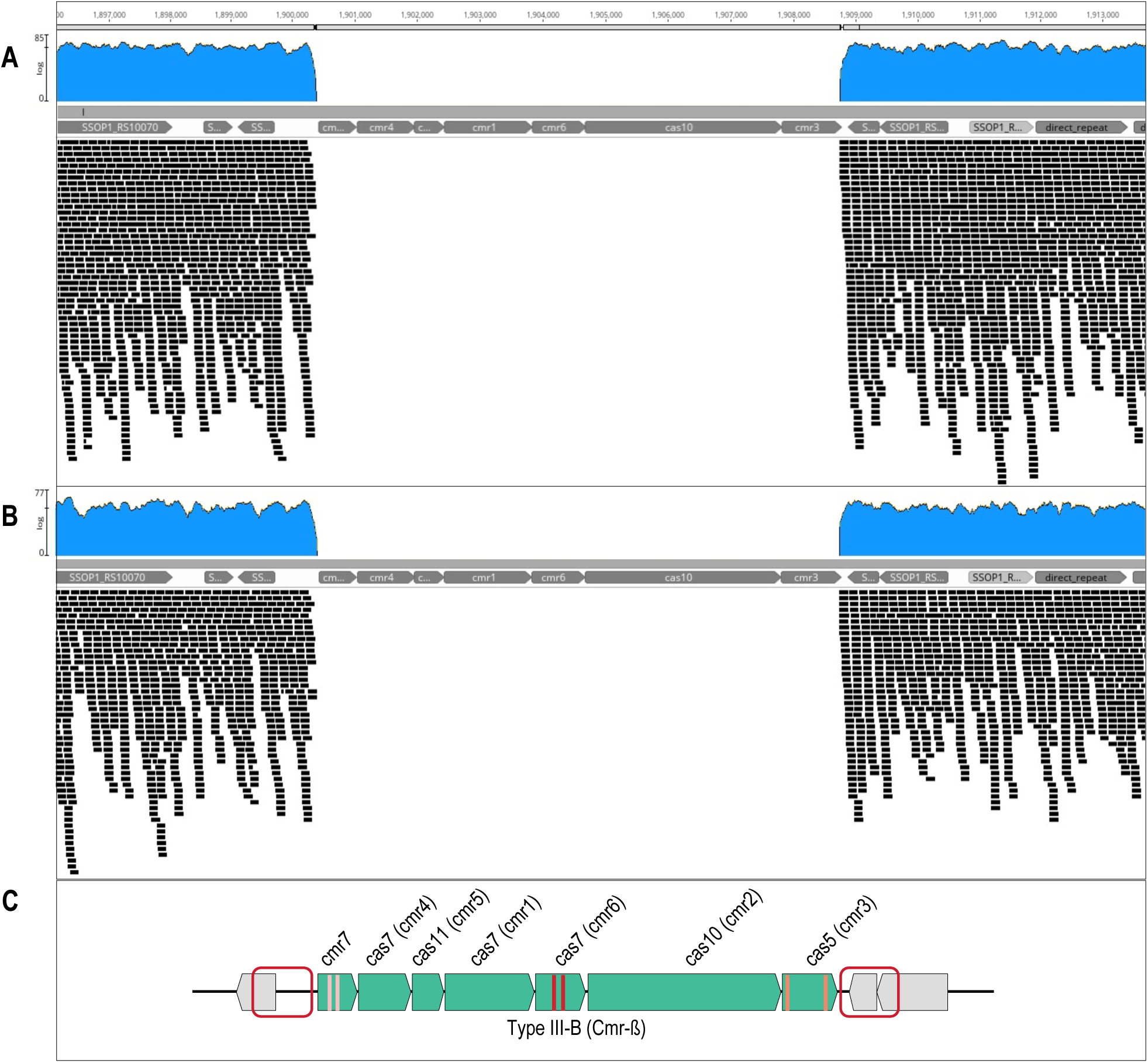
Genome viewer snapshot of *cmr-ß* knockout mutants. Paired-end reads (150 bp) generated from **(A)** KO1 and **(B)** KO2 were mapped to the *Sa. solfataricus* P1 NCBI reference genome (NZ_LT549890.1). Reference genes are displayed as grey arrows. The upper panel shows read coverage (blue) and the lower panel shows aligned reads (black). The excision of the *cmr-ß* gene cluster and its promoter as depicted in **(C)** was confirmed to have happened as intended for both isolated mutant strains.

**Figure 3.**
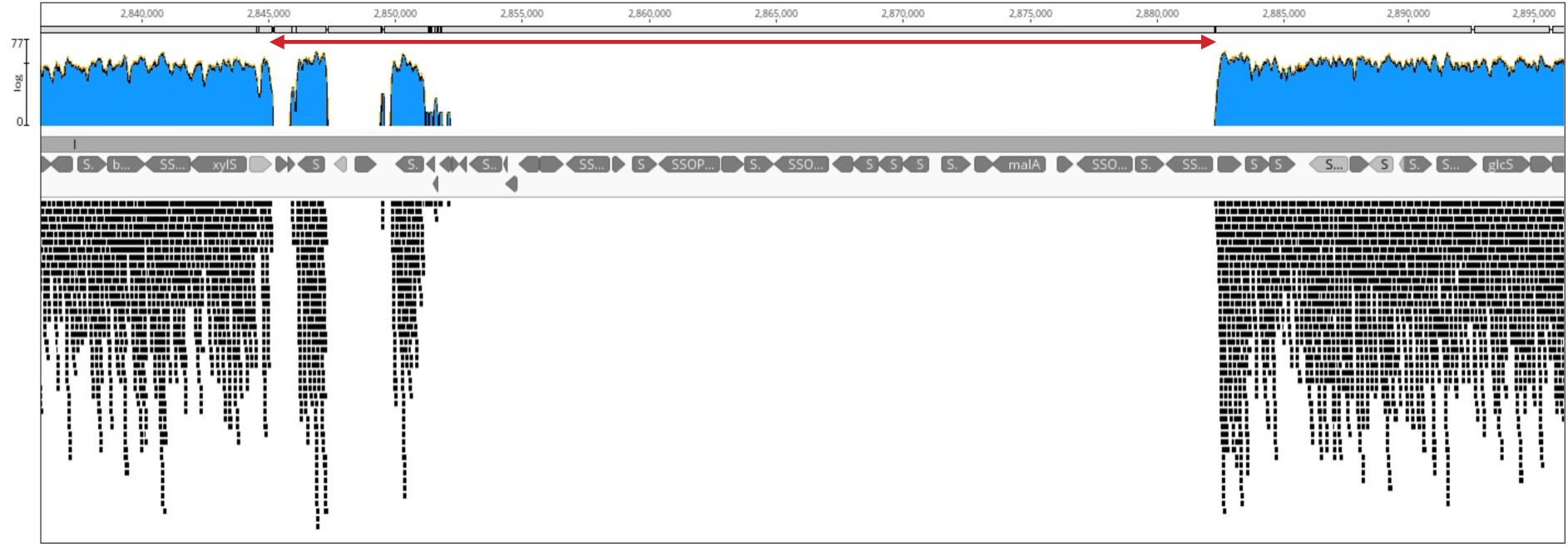
Genome viewer snapshot of second deletion that occurred spontaneously in *cmr-ß* mutant KO2. Paired-end reads (150 bp) were mapped to the *Sa. solfataricus* P1 NCBI reference genome (NZ_LT549890.1). Reference genes are displayed as grey arrows. The upper panel shows read coverage (blue) and the lower panel shows aligned reads (black). The deleted region indicated by a red arrow spans over 37 kb and is borderd by two IS5 family transposase (pseudo-) genes. Read alignments within the indicated deleted region correspond to copies of different IS4 family transposase genes.

### No difference in plasmid immunity between *cmr-ß* knockout mutants and wildtype strains

To investigate the contribution of Cmr-ß to the immune defense against foreign genetic elements in *Sa. solfataricus* P1, both wildtype and *cmr-ß* knockout mutant were challenged with two plasmids generating a transcript of a protospacer (i.e. target) complementary to a crRNA encoded in CRISPR array D (Fig. 4A and B). We have previously demonstrated that cells transfected with a viral vector carrying the same protospacer flanked by a PAM could not survive, even under selective conditions^24^, indicating an active type I CRISPR-Cas immune response. To solely study the immune response of Cmr-ß and/or Csm by circumventing type I CRISPR-mediated interference, the PAM of construct D2.109 was now disrupted by nucleotide insertion at the 3’ end of the protospacer. Upon recognition of this protospacer transcript, the available type III effector(s) should elicit a multi-layered immune response causing detrimental effects to the cells due to abortive infection. As mentioned above, abortive infection can be abolished by the presence of a sequence stretch at the 3’ end of the protospacer (antitag) complementary to the 5’ tag of the crRNA while the bound target is still degraded^7–9,11–14,37,38^. Therefore, we equipped the same protospacer with a 3’ flanking antitag in construct D2.109-a (Fig. 4A), which in our previous study yielded a type III-CRISPR-Cas-mediated reduction in protospacer transcript levels while DNA interference was not observed^24^. In addition, two different controls were used, namely cells transformed with plasmid A53 lacking a matching protospacer^24,39^ and untransformed cells not harboring any plasmid (negative control, NC) (Fig. 4A, 4B). After introduction of the different plasmids into both wildtype and deletion mutant, it became evident that growth of cells harboring D2.109 was negatively affected to a similar extent in both wildtype and *cmr-ß* knockout mutants (Fig. 4D, Suppl. Fig. S2 and S3). In both strains, cells transformed with A53 and D2.109-a were still growing in a 10fold higher dilution on plates compared to cells carrying plasmid D2.109. The same overall trend could also be observed when following their growth in liquid culture (Fig. 4C). Consequently, the absence of the Cmr-ß complex had no obvious effect on the extent of the observed type III CRISPR-Cas-mediated immune response in our experimental setting.

**Figure 4.**
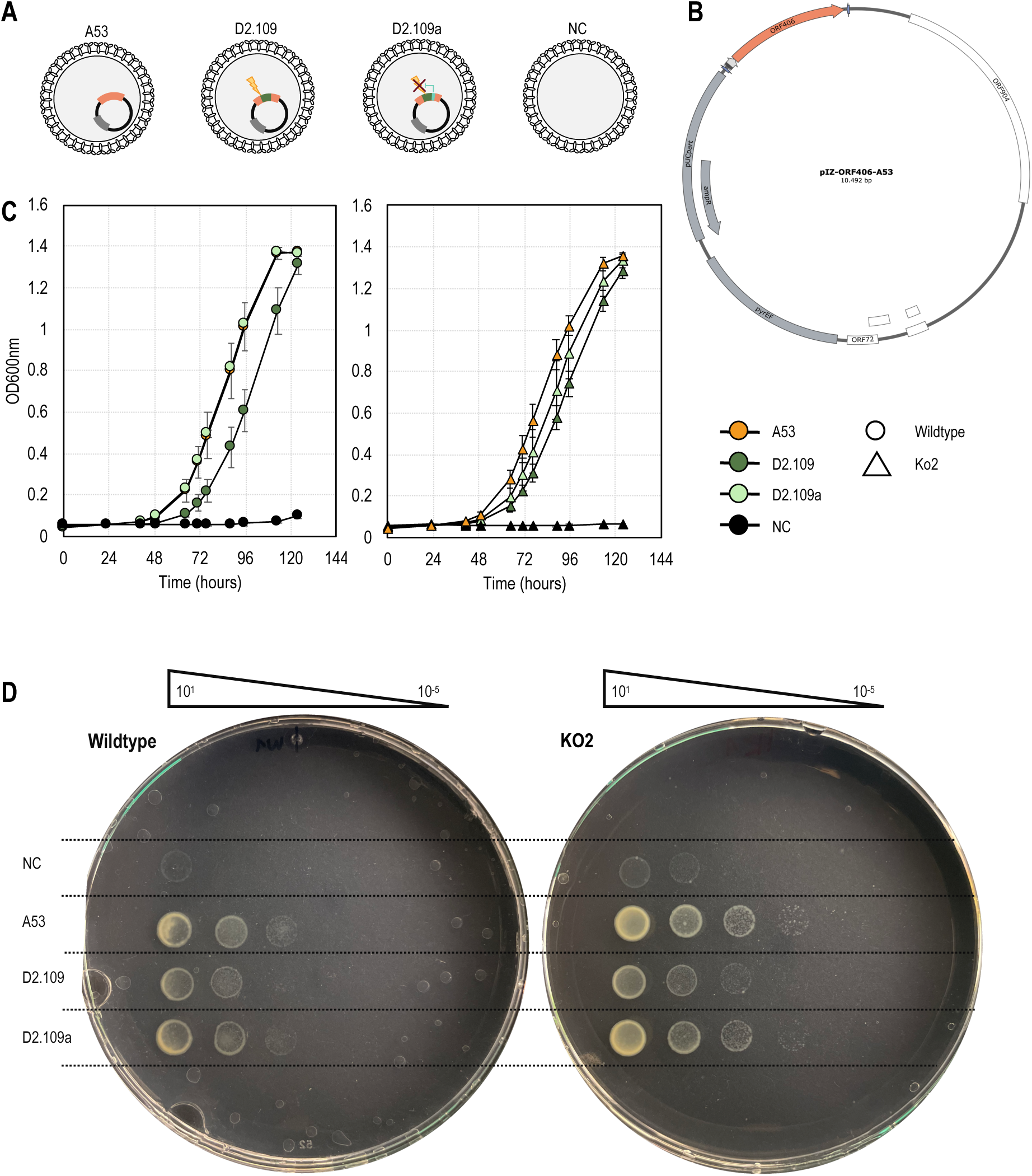
Plasmid challenge experiment. **(A)** Wildtype and deletion mutant KO2 were transformed with plasmids harboring different protospacers: A53 is not recognized by the CRISPR immune system due the lack of a matching spacer; D2.109 matches an endogenous spacer in CRISPR locus D eliciting type III CRISPR-mediated immunity; D2.109a harbors the same protospacer as in D2.109 and a matching 3’ antitag to inhibit abortive infection. Negative control (NC) cells did not receive a plasmid and therefore lack an intact *pyrEF* locus. **(B)** Plasmid map of pIZ-ORF406-A53. **(C)** Growth curves for control (left) and KO2 (right) were generated by measuring optical density at 600nm, samples are color coded as follows: NC -black; A53 - orange; D2.109 - dark green; D2.109a - light green. Error bars: mean ± SD (n = 3, i.e. three separate inoculations after electroporation). **(D)** After transformation, cells were recovered in selective media before performing 10-fold serial dilutions (10^1^ to 10^-5^) and spotted on plates. One representative plate per strain is shown, the remaining plates are provided in the supplementary material (Suppl. Fig. S2 and S3).

### Knockout mutants exhibit faster growth and significant changes in their transcriptional landscape compared to the wildtype

To test whether growth of *cmr-ß* knockout mutants was affected compared to the wildtype in complex medium, four replicate cultures for each strain were inoculated to the same initial OD_600_ (0.027 ± 5.88E-04). In comparison to the wildtype, similarly accelerated growth was observed for both knockout strains KO1 and KO2, the latter being the strain that attained the additional genomic deletion (Fig. 5A). Samples were taken at the start of exponential growth at an OD_600_ = 0.2 and RNA was extracted to perform RNA-seq. Principal component analysis (PCA) revealed a distinct shift in the transcriptomic landscape upon deletion of Cmr-ß (Fig 5B). All four replicates of each strain grouped together, thereby indicating high reproducibility. Additionally, the two deletion strains were overlapping whereas the wildtype was clearly clustering separately. One principal component (PC1) could explain ∼39% of the variance between knockout strains and wildtype.

**Figure 5.**
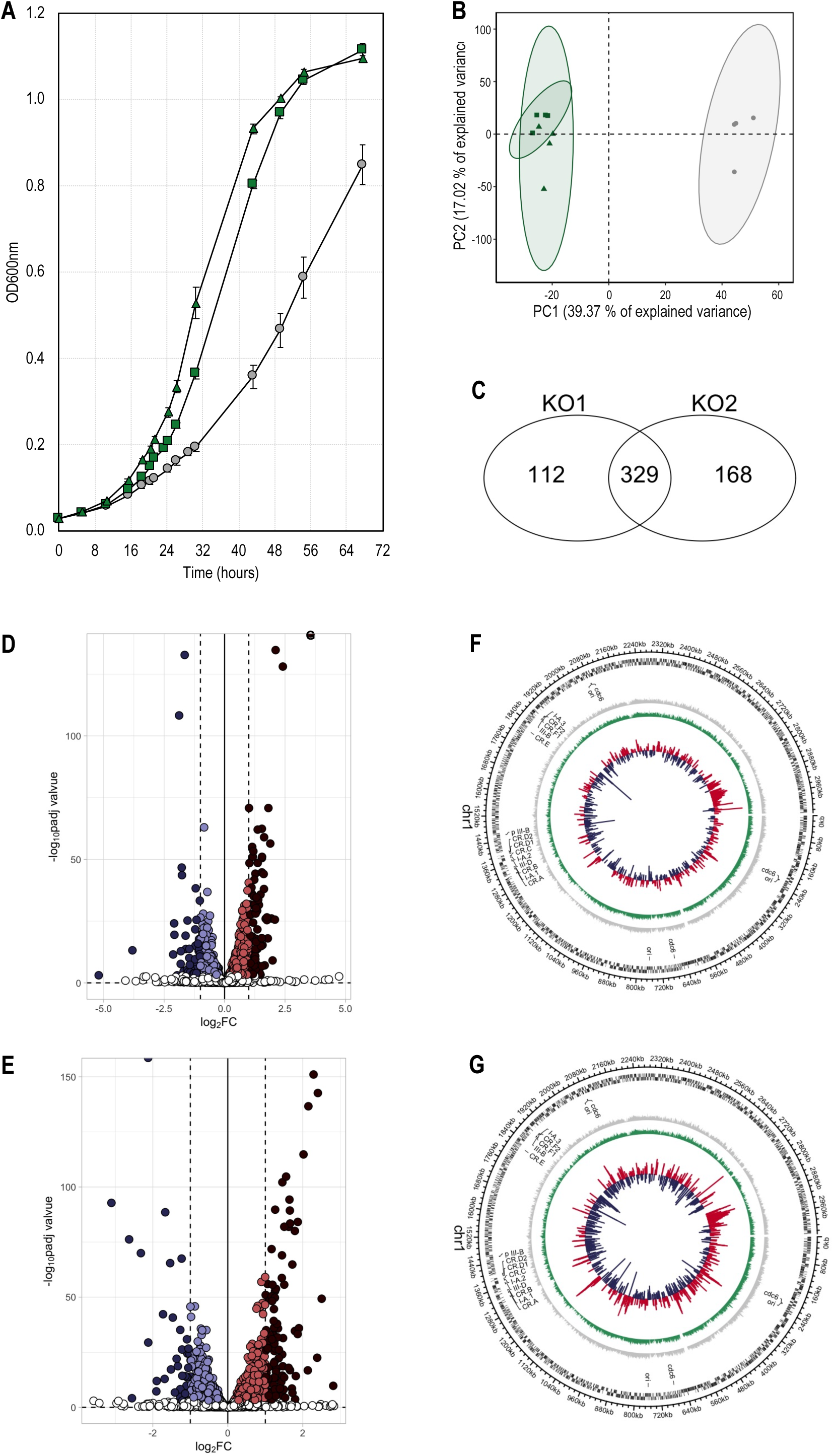
Knockout of the *cmr-ß* genes leads to growth accelaration and significant changes on the transcriptomic level. **(A)** Initial growth curve following isolation of two separate *cmr-ß* knockout mutants in comparison to wildtype undergoing the same procedure for the isolation of a single colony. Growth was followed by measuring optical density at 600nm. Color code: wildtype control – grey; deletion mutant 1 (KO1) – dark green; deletion mutant 2 (KO2) – light green. Error bars, mean ± SD (n = 4, i.e. four separate inoculations). **(B)** Principal component analysis of log transformed transcriptomic data from early exponential growth phase. Ellipses indicate 95% confidence interval. Same color code as in growth curve is used. **(C)** Venn diagram showing overlap between KO1 and KO2 regarding significantly upregulated genes in comparison to wildtype. **(D)** and **(E)** Volcano plots displaying log2-fold changes (x-axis) and associated significance (-log10 adjusted p value, y-axis), comparing KO1 and KO2, respectively. Color code indicates genes that are strongly upregulated (log2FC > 1, padj ≤ 0.001, dark red), upregulated (log2FC < 1, padj ≤ 0.001, red), strongly downregulated (log2FC < −1, padj ≤ 0.001, dark blue), downregulated (log2FC > −1, padj ≤ 0.001, blue). **(F)** and **(G)** Circular genome-wide plot for KO1 and KO2, respectively, showing *Sa. solfataricus* genome with position 0 to the right. Position of origins of replication (ori) and corresponding *cdc6* genes, as well as CRISPR-Cas systems are indicated. From outside inwards, rings indicate annotated coding sequences (black), log2-transformed TPM values for control (grey) and knockout mutant (green). Log2-fold changes in expression levels are displayed for significantly upregulated (padj ≤ 0.001, red) and downregulated (padj ≤ 0.001, blue) genes.

Changes in gene expression with high statistical significance (cutoff for adjusted *P* value (padj) < 0.001) were taken into consideration in the downstream analysis even in case of smaller log2 fold changes (FC) between deletion mutant and control. This approach is in accordance with previous transcriptomic studies in other archaea that have reported lower log2 fold changes of statistical significance for genes of physiological relevance^40–42^. Of the 3002 coding genes in *Sa. solfataricus* P1, roughly 30 % were differentially expressed in both KO1 and KO2 when compared to the wildtype (Fig. 5E and F, respectively). The majority of significantly up- and downregulated genes was shared between the two knockout mutants KO1 and KO2 (Fig. 5C and D, respectively). The top 25 upregulated genes for KO1 and KO2 are listed in Supplementary Tables S2 and S3, respectively. Observed differences were seemingly not caused by a polar effect on the up- and downstream genes surrounding the site where the targeted deletion had occurred, as significant changes in transcriptional levels of genes were dispersed throughout the genome (Fig. 5G and H). A functional enrichment analysis based on archaeal clusters of orthologous genes (arCOGs) was performed to determine which general biological functions were affected by the deletion (Fig. 6A and B). Genes involved in energy production and conversion (category C), amino acid transport and metabolism (category E), and nucleotide transport and metabolism (category F) were significantly enriched among the significantly upregulated genes in both KO1 and KO2. In KO1, genes associated with carbohydrate transport and metabolism were additionally found to be overrepresented among the significantly upregulated group of genes (Fig. 5A). In contrast, there were no arCOG categories specifically enriched among the downregulated group of genes.

**Figure 6.**
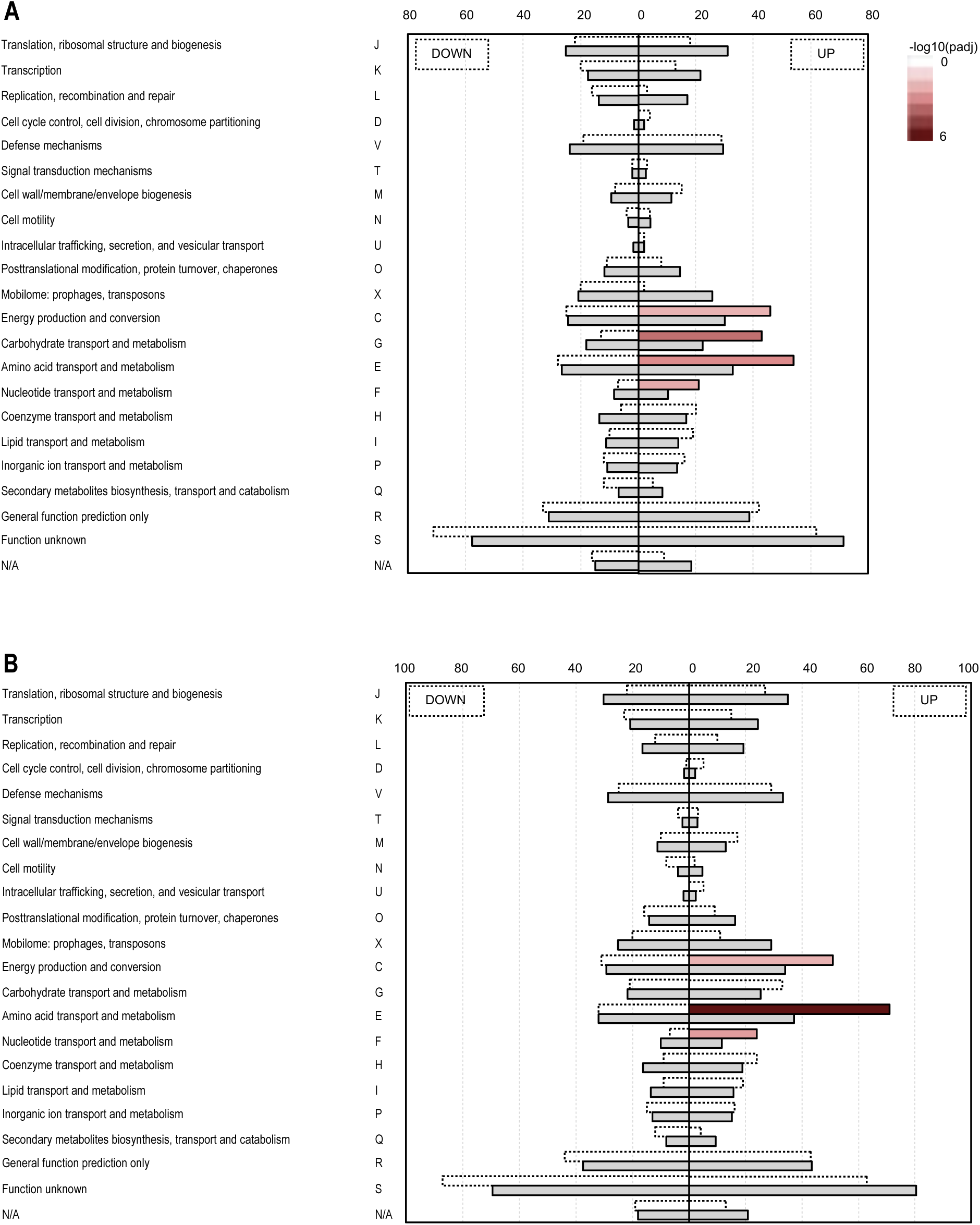
Functional enrichment analysis based archaeal clusters of orthologous genes (arCOGs) for **(A)** KO1 and **(B)** KO2. Left and right side of the plot show results for significantly downregulated and upregulated genes, respectively. Grey bar size represents the number of genes expected for each category based on a hypergeometric test, whereas neighboring bars represent the found number of genes for each category. Categories with padj-value < 0.05 are considered significantly overrepresented and highlighted by solid lines around the bar. Significance levels (-log10 transformed adjusted p-value) are indicated by color scale from white to dark red.

### Absence of Cmr-ß results in significant changes in core metabolic modules

In accordance with the functional enrichment analysis and the accelerated growth observed in *cmr-ß* knockout strains, several key components of the energy metabolism were upregulated in both knockout strains (all pathways discussed in this section are summarized in Fig. 7, Supplementary Notes). Regarding the electron transport chain (ETC), most of the genes encoding subunits of complex I (NADH dehydrogenase) were upregulated apart from NADH dehydrogenase subunit B (NuoB), which is encoded elsewhere in the genome. In Sulfolobales, c-type cytochromes are absent and the functions of complex III and IV of the ETC are combined within quinone oxidoreductase supercomplexes^43^. These unusual terminal oxidase complexes accept electrons directly from quinones, then in turn reduce molecular oxygen and pump protons to generate a proton motive force. Transcription levels of two out of three terminal oxidases in *Sa. solfataricus* were increased in *cmr-ß* knockout strains. While all genes encoding for terminal oxidase SoxEFGHIM were upregulated, only half of the genes encoding for terminal oxidase SoxABCDL showed significantly higher expression. The hydrogenase-related subunits of a putative formate hydrogenlyase complex (FHL) present in the genome were significantly upregulated in knockout strains, while the expression levels of the putative formate dehydrogenase subunit (FdhA, a molybdenum-dependent enzyme) were unchanged, or very slightly downregulated. In accordance with that, the periplasmic component of the ABC molybdate transport system located downstream of the FdhA was significantly downregulated.

**Figure 7.**
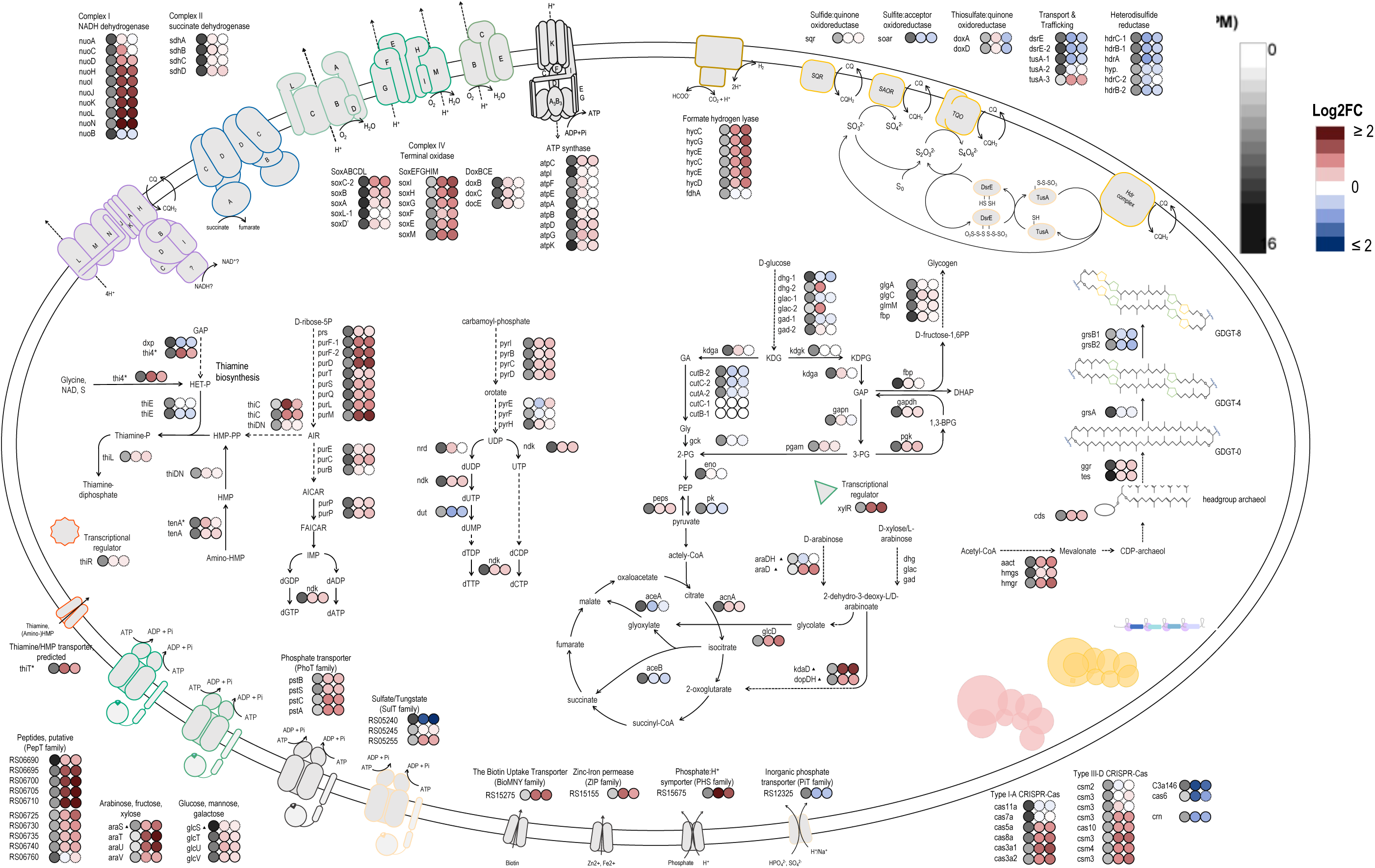
Response in metabolism to deletion of type III-B CRISPR-Cas system. First circle from left to right represents log2-transformed TPM values as an approximation for expression levels in wildtype control, indicated by color bar from white (not expressed) to black (highly expressed). Log2FC in transcript levels found in KO1 and KO2 are represented in that order by the two following circles, indicated by color scale from dark blue (strongly downregulated) to dark red (strongly upregulated). Significant change (padj ≤ 0.001) is indicated by solid outlines around circles. Genes with predicted binding sites for transcription factors ThiR or XylR are indicated by star or triangle symbols, respectively.

Another process that was seemingly impacted on a transcriptional level by the *cmr-ß* knockout was the transport of nutrients, with several different types of transporters being significantly upregulated. Regarding organic carbon compounds, the arabinose ABC transporter (3.A.1.1.14-AraS) and several putative peptide ABC transporters (3.A.1.5-PepT) were clearly upregulated in KO1 and KO2, while the glucose ABC transporter (3.A.1.1.13-GlcS) exhibited mildly increased transcript levels^44,45^. The gene encoding transcriptional regulator XylR (SSOP1_RS15310), which in *S. acidocaldarius* was shown to positively regulate the expression of pentose transporters as well as metabolizing genes^46^, was significantly upregulated. Accordingly, all but one gene with the corresponding predicted binding site for XylR^47^ were upregulated, among them *araS* and *glcS* encoding the substrate-binding proteins of the respective ABC-transporters (Fig. 7). Interestingly, some genes of the XylR regulon were recently reported to be part of an independently modulated set of genes that also included the *soxEFGHIM* gene cluster, indicating that their expression might be governed by a shared unknown regulator^48^. Transporters putatively involved in the uptake of amino acids (2.A.1.15-The Aromatic Acid:H+ Symporter Family, 2.A.3.11-The Aspartate/Glutamate Transporter Family) were also highly upregulated in both mutants.

A predicted thiamine transporter^49^ (ThiT) was highly upregulated in *cmr-ß* knockout strains, as well as at least one homolog of ThiC (HMP phosphate (HMP-P) synthase) catalyzing the first step in the synthesis of the pyrimidine moiety of thiamine (HMP-PP), and Thi4 (thiamine thiazole synthase) catalyzing the formation of the thiazole ring (HET-P)^50,51^. Additionally, transcript levels of at least one out of two TenA homologs (aminopyrimidine aminohydrolase) implicated in the salvage of thiamine precursors was slightly higher in both mutant strains compared to wildtype. These genes are part of a proposed thiamine regulon in *Sa. solfataricus*, which includes genes involved in the biosynthesis pathway and transport of thiamine and/or precursors^49^.

*Sa. solfataricus* employs a reversed monophosphate pathway for the synthesis of ribose-5-phosphate, which serves as pentose moiety in nucleotide biosynthesis^52^. The genes encoding the enzymes partaking in purine de novo biosynthesis until the formation of the intermediate 5′-phosphoribosyl-5-aminoimidazole (aminoimidazole ribotide, AIR) were upregulated in KO1 and KO2. Interestingly, this intermediate compound is used as a precursor during the generation of HMP-PP, establishing a link to the observations regarding thiamine biosynthesis^49^. In addition, slightly increased expression levels could also be observed for enzymes participating in purine salvage. In contrast, the genes encoding enzymes that catalyze the reaction from cytosine to uracil in the process of pyrimidine salvage were significantly downregulated, likely due to the supplementation of uracil in the medium.

Genes encoding TatC (SSOP1_RS15345) and SecY (SSOP1_RS03615), components of two different protein translocation pathways^53–56^, were also significantly upregulated in *cmr-ß* knockout strains. Interestingly, insertion of TatC itself into the membrane was shown to require the Sec pathway together with translocase YidC in E. coli^57^, which we also found significantly upregulated in our dataset (SSOP1_RS17510). Overall, a significant enrichment of genes encoding transmembrane domain proteins could be observed among the upregulated fraction of the transcriptomes of both KO1 and KO2 (Fig. 8A and B, respectively).

**Figure 8.**
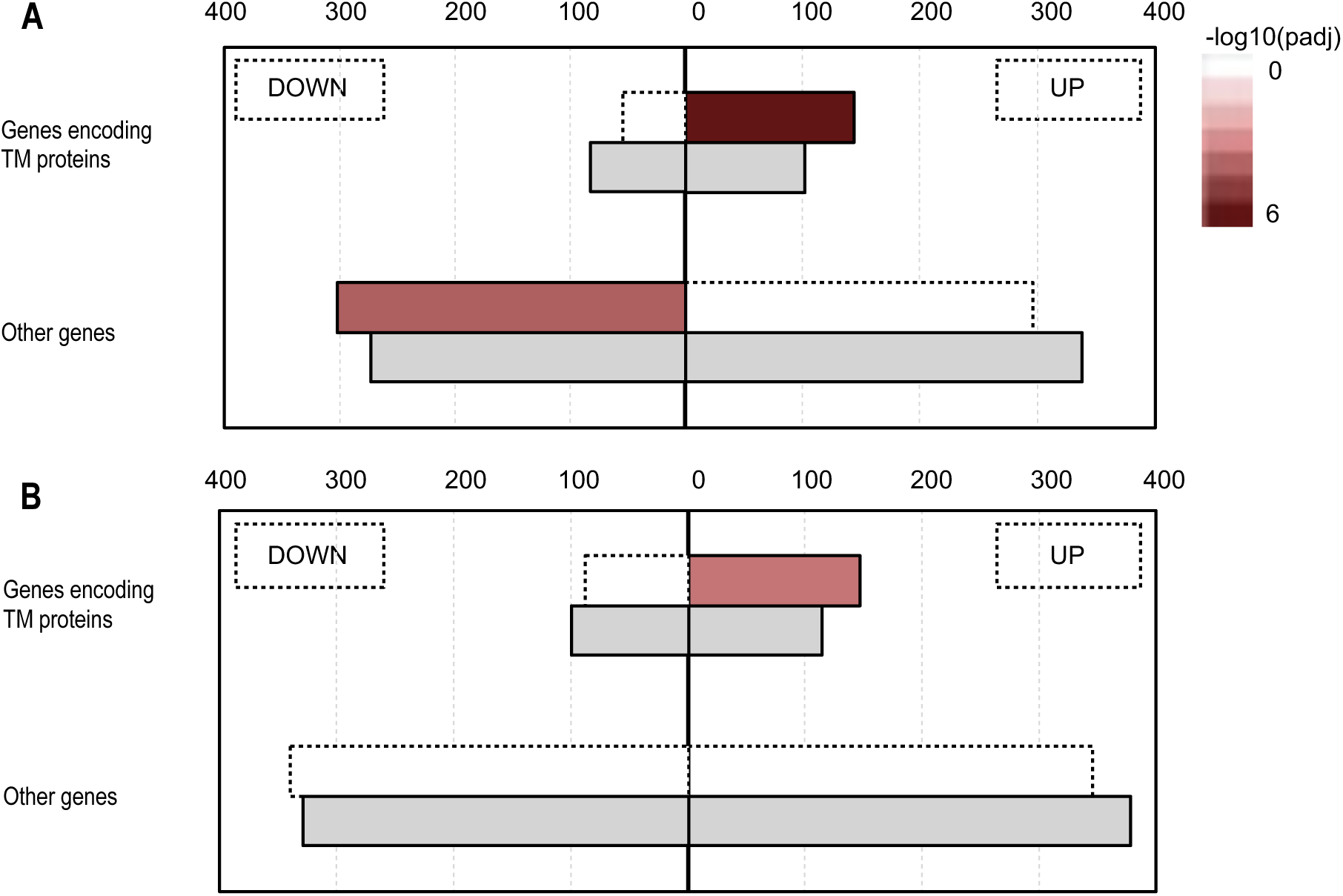
Enrichment analysis of genes encoding transmembrane-domain proteins as predicted for protein products by phobius. for **(A)** KO1 and **(B)** KO2. Left and right side of the plot show results for significantly downregulated and upregulated genes, respectively. Grey bar size represents the number of genes expected for each category based on a hypergeometric test, whereas neighboring bars represent the found number of genes. Categories with padj-value < 0.05 are considered significantly overrepresented and highlighted by solid lines around the bar. Significance levels (-log10 transformed adjusted p-value) are indicated by color bar from white to dark red.

A number of genes involved in the isoprenoid lipid biosynthesis were significantly upregulated in the *cmr-ß* knockout strains. Specifically, the genes responsible for the synthesis of intermediates mevalonate (thiolase -*aact*, hydroxymethylglutaryl-CoA synthase - *hmgs*, hydroxymethylglutaryl-CoA reductase - *hmgr*), glycerol-1-phosphate (glycerol-1-phosphate dehydrogenase) and the ether bond formation step to geranylgeranylglyceryl phosphate (GGGP) synthase), as well as geranylgeranyl reductase (*ggr*) and tetraether synthase (*tes*), leading to the formation of the membrane-spanning glycerol dibiphytanyl glycerol tetraethers (GDGTs). Interestingly, the modifying enzyme calditol synthase (*cds*) responsible for the synthesis of the cyclopentenyl head group calditol^58^ was significantly upregulated, while GDGT ring synthases *grsA*, *grsB1* and *grsB2*^59^ were downregulated, the latter two significantly.

Lastly, 28-32 putative transcriptional regulators (arCOG functional category K), characterized and uncharacterized, were significantly differentially expressed in the knockout strains, indicating a strong involvement of the Cmr-ß complex to various regulatory circuits. However, the CRISPR-associated Csa3 transcription factors involved in regulation of various CRISPR-Cas system components and coordinating the immune response^60–64^ were not differentially expressed according to our cutoffs.

### Response of CRISPR-Cas defense systems and transposases to the deletion of Cmr-ß

Next, we wanted to investigate whether the absence of Cmr-ß had influenced the expression of other genes implicated in antiviral defense. Looking at the expression levels of CRISPR-*cas* modules in our wildtype samples, it became evident that the first out of three type I-A *cas* modules (SSOP1_RS07365-SSOP1_RS07390) as well as the *cmr-ß* module were most highly expressed. The second and third type I-A module were considerably lower and basically not expressed, respectively. In comparison, the expression level of the *csm* module was more than ten times lower than the one of the *cmr-ß* module under standard growth conditions. Upon *cmr-ß* knockout, we could detect an upregulation of the majority of genes of the first type I-A *cas* module as well as increased expression levels for the *csm* module (Fig. 7).

Several transposase genes were significantly downregulated in absence of Cmr-ß. The genus *Saccharolobus* generally carries a remarkable number of insertion sequence (IS) elements^65–68^. For *Sa. solfataricus* P2 specifically it is estimated that IS elements make up approximately 10% of its total genome sequence (in bp)^69^. In the P1 reference genome, almost 170 functional transposases are currently annotated (based on eggnog annotations of NZ_LT549890.1). Several transposases of the most prominent IS4 (ISC1225, ISC1359, ISC1439) and IS5 (ISC1058, ISC1212, ISC1234) families were found to be downregulated in both deletion mutants. Transposase genes belonging to the less abundant IS630 (ISC1048, ISC1078), ISC1217 and IS110 (ISC1190, ISC1229) elements were also exhibiting lower transcript levels in absence of the Cmr-ß complex. In contrast, although some transposases of the IS200/IS605 and IS607 families were found to be downregulated, certain representatives were upregulated (especially in KO2).

### Adaptation to minimal media is affected in *cmr-ß* knockout mutant

Since we observed several genes related to transport being differentially expressed in complex (sucrose-tryptone) media, we were interested in examining to which extent this phenotype would still be observable in defined media. Wildtype and KO2 strain were adapted to grow in media containing either glucose or sucrose as sole carbon source. The switch from complex to defined media caused a lag phase in both strains, however it was more pronounced in the *cmr-ß* knockout strain (Fig. 9A). The switch to sucrose resulted in a lag phase of about 48 hours (hr) for the wildtype strain, after which it assumed a growth rate similar to the one exhibited under complex media and reached stationary after 100 hr, while the knockout endured both a prolonged lag phase of about 72-96 hr as well as an overall reduced growth rate with linear growth for the next 96 hr, reaching stationary phase after 192 hr. The switch to glucose media proved more cumbersome for both strains and resulted a slightly different profile. The wildtype endured a lag phase of about 72 hours, after which it followed linear growth for the next 138 hours, while the knockout exhibited a very long lag phase of 264 hr, after which it showed exponential growth with a growth rate only slightly lower than in complex media. After adaptation, the growth curves of wildtype and KO2 ultimately converged in glucose medium, but not in sucrose, where the wildtype grew slower than the knockout (Fig. 9B and C, respectively). Of the differentially expressed genes in glucose medium, 234 (∼8 % of protein-coding genes) were upregulated and 260 (∼9 % of protein-coding genes) were downregulated in KO2 (not shown). In sucrose medium, more than twice as many genes were found to be significantly differentially expressed in KO2, among those 621 (∼20 %) were upregulated and 560 (∼19 %) were downregulated (not shown). The complex media condition served as an internal control in this second experiment, and we found 287 (∼10 %) and 254 (∼9 %) genes to be up- and downregulated in KO2, respectively (not shown). Here, the vast majority of upregulated genes was consistent with our previous dataset (∼73 %) (Fig 10A and Supplementary Information). Interestingly, 20 genes were similarly upregulated in all three conditions according to DESeq analysis (Suppl. Tab. S4). Together with genes exhibiting a strong correlation with all *cmr-ß* knockout samples in combined PCA analyses (Fig. 10B, Suppl. Tab. S5), the overarching theme again points towards perturbations in energy metabolism, transport and lipid biosynthesis following the *cmr-ß* knockout. The higher expression of these genes, regardless of medium composition, strongly implicate a role of the Cmr-ß in the regulation of metabolism either directly or indirectly.

**Figure 9.**
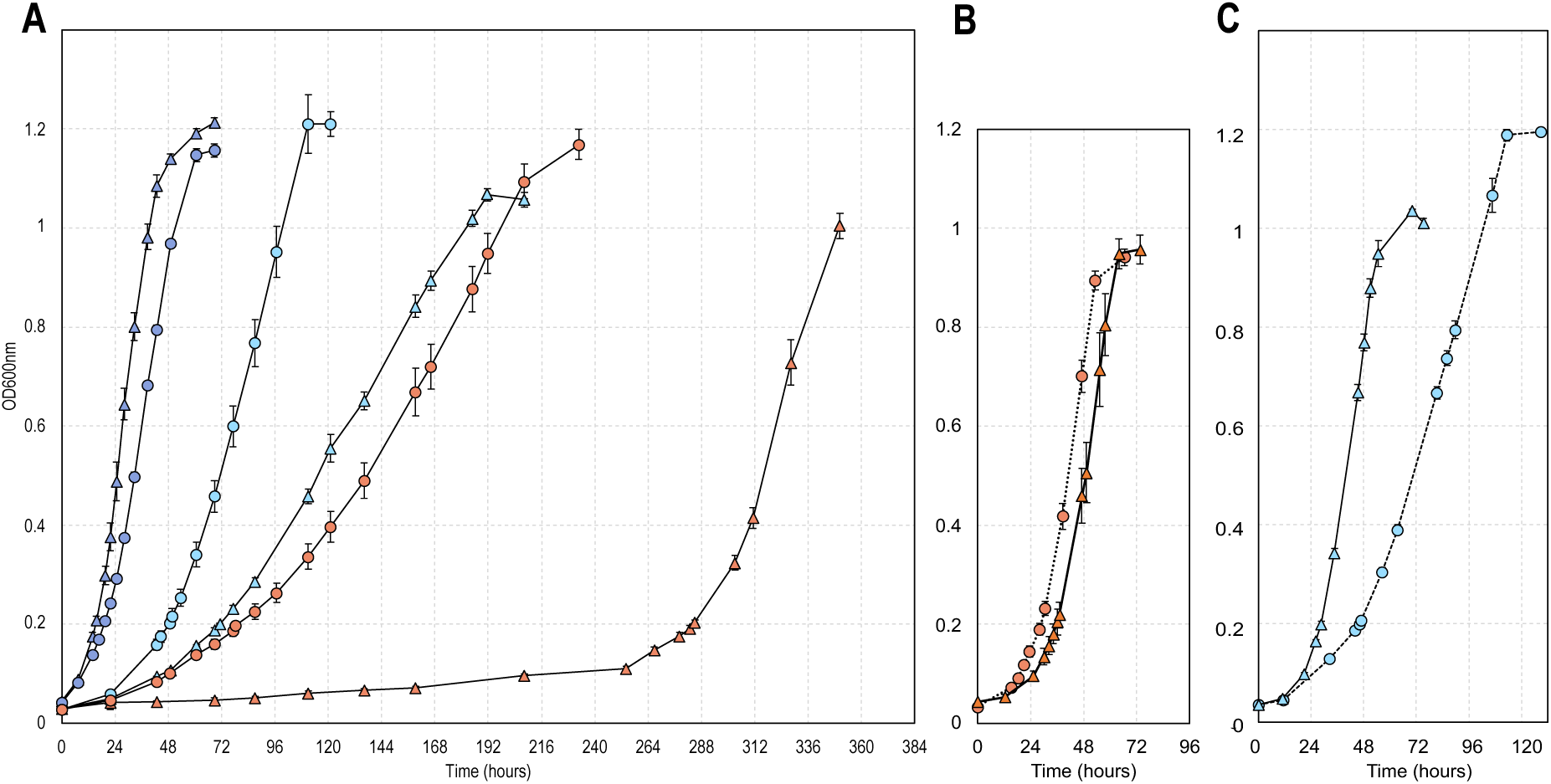
Adaptation Experiment Growth Curves. **(A)** Switching wildtype control and deletion mutant from complex to defined carbon sources sucrose and glucose causes lag phase during adaptation. Growth was followed by measuring optical density at 600nm. Wildtype control and Ko2 samples are represented by circles and triangles, respectively. Samples are color coded according to media composition as follows: Sucrose-tryptone – blue; sucrose – light blue; glucose – orange. Error bars, mean ± SD (n = 4, i.e. four separate inoculations). **(B)** and **(C)** show growth of KO2 and wildtype in glucose and sucrose media, respectively, after adaptation.

**Figure 10.**
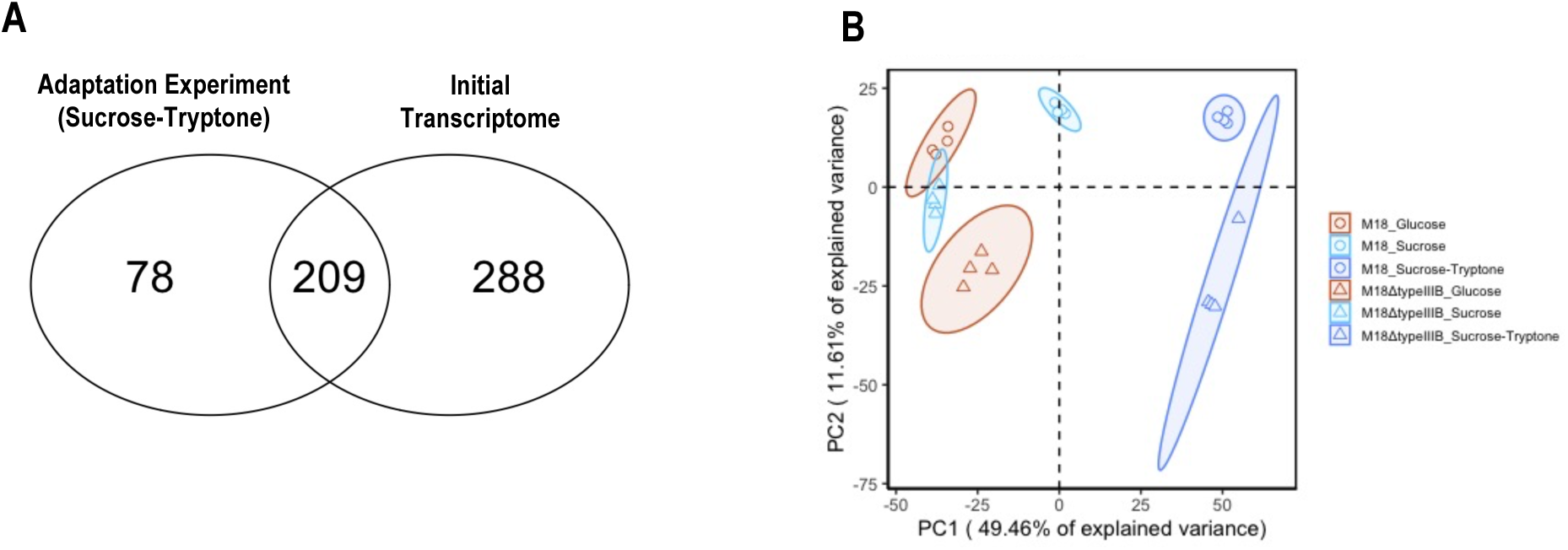
Transcriptomic Analysis of Cultures Adapted to Glucose and Sucrose Medium. **(A)** Venn diagram showing overlap in significantly upregulated genes between initial transcriptomic dataset and sucrose-tryptone condition of adaptation experiment. **(B)** Principal component analysis of log transformed transcriptomic data from early exponential growth phase. Ellipses indicate 95% confidence interval. Wildtype control and KO2 samples are represented by circles and triangles, respectively. Samples are color coded according to media composition as follows: Sucrose-tryptone – blue; sucrose – light blue; glucose – orange.

## DISCUSSION

### Cmr-ß might not be a major contributor to CRISPR-Cas-mediated plasmid defense in *Sa. solfataricus*

The co-existence of two different type III CRISPR-Cas complexes in *Sa. solfataricus* and other Sulfolobales^70^ might seem redundant at first, however is most likely indicative of different mechanistic specialization which enables complementation in response to varying load and types of MGEs^71,72^. In this study, we set out to investigate the specific roles of Cmr-ß in the immune response of *Sa. solfataricus* P1. Interestingly, while *cmr-ß* knockout mutants were readily obtained, similar attempts at deleting the *csm* locus caused large genomic deletions (Wimmer unpublished) and isolated mutants could therefore not be tested. Our plasmid challenge assay indicated that the transcription-dependent targeting effect of a protospacer-carrying plasmid in *cmr-ß* knockout mutant was comparable to the wildtype. Hence, *Sa. solfataricus* P1 Csm-mediated immunity leads to a similar outcome irrespective of the presence of the Cmr-ß system (see Fig. 4). In analogy to SisCmr-ß^33^, one could assume that Cmr-ß of *Sa. solfataricus* P1 does not synthesize significant amounts of cOA molecules and is possibly not a major contributor to the canonical multi-faceted type III CRISPR -mediated defense.

### Transcriptional response to the *cmr-ß* knockout supports accelerated growth phenotype and interaction with regulatory networks

The findings of our transcriptomic analysis show that the deletion of Cmr-ß had significant effects on the transcriptional landscape and growth behavior. In accordance with accelerated growth following the deletion, we could observe upregulation of genes involved in distinct functional categories of energy production and conversion, as well as metabolism and transport of amino acids, nucleotides, and carbohydrates. The upregulation of complex I (NADH dehydrogenase) and two out of three complex IVs is interesting regarding their importance in maintaining pH homeostasis and generating a proton motif force in thermoacidophilic organisms^43^. This observation is even more compelling considering that complex I is capable of pumping protons as opposed to complex II (succinate dehydrogenase)^73^ and the bb3-type terminal oxidase has a higher electron to proton pumping ratio than the other terminal oxidases (aa3-type SoxABCDL and DoxBCE) due to the presence of two proton pumping sites^74,75^. Moreover, its subunit SoxE was suggested to act as a pH sensor and regulate electron flow^74,76^. The activity of characterized FHL complexes has also been connected to counteracting cytoplasm acidification through removal of formate and H2 evolution^77,78^. Moreover, being membrane-bound complexes with similarities to complex I, it has long been hypothesized that they might be capable of proton pumping and therefore contribute to proton motive force generation, a capacity strongly supported by the formate-driven growth in *Thermococcus onnurineus*^79,80^. Both these activities might explain the upregulation profile of the hydrogenase component of the FHL in the knockout strains.

Upregulation of ABC-type and other transporters for the uptake of substrates such as sugars, peptides/amino acids and phosphate further supports the accelerated growth phenotype of the knockout strains. The additional upregulation of thiamine transport and biosynthesis as well as nucleotide biosynthesis seemingly underlines the need for higher ATP synthesis and energy generation. The increased incorporation of membrane-anchored proteins could be supported by the upregulation of pivotal components of protein translocation pathways, namely *tatC, secY* and *yidC*. As central component of the Tat pathway, TatC captures folded proteins destined for translocation such as subunits of terminal oxidases that assemble their cofactors prior to translocation in the cytoplasm^81,82^. In contrast, ABC transporters of CUT class of *Sa. solfataricus* harbor type IV pilin like signal peptides and are presumed to be secreted via the Sec pathway^44,45,83^.

As summarized above, the upregulation of the ETC complexes and the FHL, as well as the increased organic carbon availability implied by the upregulation of the respective transporters supports the increase in growth rate observed in the deletion mutants under complex media. The observed upregulation in main lipid metabolism genes as well as downregulation of the ring synthases would also agree with this phenotype, as it would be the expected profile under logarithmic growth without energy limitations^84–87^. The observed profile in the *cmr-ß* knockout could also indicate that *Sa. solfataricus* is experiencing (or displaying the transcriptional profile of) a mild case of acid stress, as suggested by the upregulation of calditol synthase^58^. The general transcriptional profile however is not reminiscent of a classic case of acid stress, as documented in *Sa. islandicus*, which displayed decreased energy production, upregulation of the proton-pumping ATPase, upregulation of *grsB*, upregulation of repair and turnover pathways and increased anabolism^88^. As always though, transcriptomic profiles need to be interpreted with caution, as it has been shown that the observed profiles do not always translate into matching proteomic or lipidomic profiles^88^. An unclear profile in our case is nevertheless not surprising, as the actual imposed condition (the absence of the *cmr-ß* module) is neither an environmental stressor nor a manipulation of nutrient levels, and therefore the observed phenotypic changes cannot be interpreted in a strict framework. Rather, we are getting a glimpse of the regulatory networks and processes that the Cmr-ß is involved in in the absence of external stimulants, thereby sometimes leading to contradictory profiles.

### Functional link between CRISPR systems and cell envelope processes

As outlined above, our transcriptomic analysis of type III-B CRISPR-Cas deletion mutants repeatedly pointed towards an involvement with membrane-associated processes. Accordingly, genes encoding for transmembrane proteins were significantly enriched among the upregulated fraction in type III-B deletion mutants under all conditions examined (Fig. 8). In general, a connection between CRISPR-Cas systems and membrane-associated processes seems intuitive as it would allow for quick responses upon entry of infecting viruses.

First hints at the functional link between the cell envelope and CRISPR-Cas were already gathered relatively early on, when it was shown that the incorporation of misfolded proteins translocated by the Tat system likely caused proton leakage at the membrane prompting the activation of the type I CRISPR-Cas system in *E.coli*^89^. In the human pathogen *Francisella novicida*, Cas9 complexed with a small CRISPR-associated RNA (scaRNA) was found to transcriptionally repress Blp, a membrane-exposed lipoprotein usually recognized by the hosts immune cells^90,91^. In the gram-negative bacterium *Serratia*, the Rcs signaling pathway involved in outer membrane stress response^92^ downregulates type I and type III CRISPR-Cas defense while simultaneously enhancing innate immunity, e.g through repression of viral cell-surface receptors^93^. Genome analyses have also pointed out the enriched presence of various predicted membrane proteins in the genomic neighborhood of type III CRISPR-Cas systems, indicating a close association with the cell envelope^27,28^, although the exact nature of this proposed functional connections largely remains to be investigated *in vivo*.

### The metabolic burden of maintaining CRISPR-Cas systems

Some of the observed changes in the transcriptional landscape upon deletion of the *cmr-ß* gene cluster could also be explained by the reduced metabolic burden and cost associated with the maintenance of defense systems. In *Streptococcus thermophilus* DGCC7710 harboring multiple CRISPR-Cas systems (2 x type II, type I and type III), the disruption of one of the *cas9* genes gives mutants a competitive advantage in competition with wildtype cells, possibly due to a reduced lag phase^94^. In contrast, *Pseudomonas aeruginosa* PA14 possesses only one type I-F CRISPR-Cas system, and its deletion caused reduced comparative fitness in competition assays^95^. In general, the studies in *P. aeruginosa* suggest that constitutive fitness costs associated with the presence of functional CRISPR-Cas systems are low, however conditional costs arise due to the elicitation of the immune response upon MGE exposure^95–97^. Similar to what has been observed for *S. thermophilus*^94^, the deletion of one out of several co-existing *cas* gene modules also caused an accelerated growth phenotype in *Sa. solfataricus* P1. However, the significant congruence between transcriptomes from two different deletion mutant strains despite the presence of a second large genomic deletion in only one of them suggests that we are, at least in part, witnessing valid functional connections with certain degree of specificity.

### Are the observed changes the result of direct regulation by Cmr-ß or secondary effects?

The observed profiles of anabolic pathways and membrane composition could be interpreted either as direct regulatory targets or indirectly affected due to the increase in available energy and resulting increased growth rate. The amount of (both characterized and uncharacterized) differentially regulated transcription factors in our dataset also indicates the entanglement of CRISPR-Cas systems with global regulatory networks. On this note, a LysR-type transcriptional regulator involved in control of secondary metabolites and motility was shown to repress type III-A immunity in the gram-negative bacterium *Serratia*^98^. Moreover, the observed differential expression of transposase genes in response to the Cmr-ß deletion could also influence host gene expression and are a known source of various ncRNAs in *Sa. solfataricus*^99–101^ and other archaea^102–106^. Former studies analyzing the content of isolated Cmr-ß and Csm protein complexes have revealed that both complexes are sequestering a proportion non-canonical guides in addition to canonical crRNAs *in vivo*^32,107^, some of which could match potential cellular targets, providing the possibility for direct CRISPR-mediated regulation. However, we also expect that many effects are downstream events of primary direct key regulation targets such as transcription factors, or reaction to the physiological changes invoked by presumed direct effects (e.g. direct upregulation of ETC transcripts would result in a change in the energy budget of the cell). Our current experimental design does not allow for an investigation into the molecular mechanisms by which the Cmr-ß could convey the observed regulatory effects, but we are working on elucidating the direct and indirect effects in the future.

## CONCLUSION

The deletion of the type III system consistently leads to an upregulation of genes encoding for membrane proteins and membrane-associated processes, the most important of which concern energy metabolism, the establishment of a proton motive force and essential nutrient acquisition, accompanied by an accelerated growth phenotype. As these results are in the absence of MGEs, we conclude that the type III system is exerting an influence on the greater metabolic functions, and therefore fitness, of the organism irrespective of its role in cellular defense.

## ACKNOWLEDGEMENTS

This research was funded in part by the Austrian Science Fund (FWF) [J 4611-B]. We thank John van der Oost, Raymond Staals, Sonja V. Albers and Xu Peng for scientific discussions.

## DATA AVAILABILITY

All datasets and supplementary tables will be made available upon publication of the manuscript.

## MATERIALS & METHODS

### Design of genetic constructs and cloning

The synthetic mini CRISPR arrays (miniCRs) containing one to three targeting spacers were assembled and cloned into pENTRY vectors as described previously^35^. Two 800 to 900 bp sequences homologous to the upstream and downstream regions of the *cmr-ß* locus were introduced into each vector to direct DNA repair mechanisms. Flank 1 corresponding to the upstream region was amplified using primers flank1_fw_SalI and flank1_rv, flank 2 corresponding to the downstream region was amplified using primers flank2_fw and flank2_rv_EagI. The sequences of all primers used for construction of genome editing constructs can be found in Suppl. Tab. S6. The resulting PCR products were checked by gel-electrophoresis and purified (Monarch PCR&DNA Cleanup Kit, NEB) according to manufacturer’s instructions. Flanks were phosphorylated using the T4 Polynucleotide Kinase (NEB) and cleaved with corresponding restriction enzyme SalI-HF or EagI-HF (NEB). pENTRY-miniCR vectors were double digested with SalI-HF and EagI-HF (NEB). Linearized pENTRY-miniCR vectors and both flanks were fused using the Quick Ligation™ Kit (NEB). Correct sequences of miniCRs and flanks were verified by Sanger sequencing and consequently transferred from pENTRY vectors into destination vector pIZ-GW^36,108^ using the Gateway™ LR Clonase™ II Enzyme mix (Invitrogen™).

### Culturing, transformation and isolation of knockout mutants

Uracil-auxotrophic derivate M18 of *Sa. solfataricus* P1 (DSM 1616, ATCC 35091) was cultivated in Brock salts medium supplemented with D(+)-Sucrose (0.2 %, w/v) (Serva), Tryptone (0.1%, w/v) (Roth) and Uracil (0.0125 mg/mL) (Sigma Aldrich), in short Brock medium S/T/U. Media pH was initially adjusted to 3.5 with H_2_SO_4_ and cultures were incubated under shaking conditions at 77 °C (Zebec 2016). Growth was assessed by measuring optical density (OD) at a wavelength of 600nm in a spectrophotometer. Competent cells for electroporation were prepared as described previously^109^. Per construct, 50 μl of M18 competent cells were mixed with 150 ng of plasmid DNA and electroporated using 1-mm cuvettes in a Gene Pulser Xcell (Biorad) with the following settings: 1240 V, 1000 Ω, 25 μF. Electroporated cells were transferred to equal volumes of recovery solution^110^ and incubated for 1 hour at 77 °C. After recovery, electroporated cells were inoculated directly into 50mL selective medium containing D(+)- Sucrose (0.2%, w/v) and NZ-Amine (0.1%, w/v) (Fluka Analytics). For plasmid counterselection, cells harboring pIZ-cmr73ts were transferred to medium containing Sucrose, Tryptone, Uracil and 5-Fluoroorotic acid (0.05 mg/mL) (5-FOA, Sigma Aldrich). Consequently, dilution streaks on gellan gum plates (0.64% w/v) (Gelrite, Roth) with the same media composition were performed. Deletion mutant colonies finally underwent three consecutive rounds of plating (without 5-FOA) to ensure the presence of a single genotype. As a control, a purified isolate of untransformed *Sa. solfataricus* M18 (wildtype) was generated using the same approach. Plates were incubated under high humidity at 77°C.

### Culture PCR

The presence of the *cmr-ß* genes was checked by adding 2 μl aliquots from OD samples or colony suspensions as template in PCR reaction with primer pairs cmr-KO-check_fw/rv or cmr-middle_fw/cmr-KO-check_rv. Successful removal of plasmid and persistence of IS-element conferring uracil auxotrophy were verified using primers ORF904_fw/rv and pyrBE_fw/rv, respectively. All primers used for culture PCR are listed in Suppl. Tab S7. For all culture PCRs, full reaction volumes were loaded on 1% agarose gels together with 1 kb Plus DNA ladder (Thermo Scientific). Knockout of *cmr-ß* indicated by smaller bands in gel-electrophoresis of PCR products was verified by sequencing of DNA from gel-purified bands (Monarch DNA extraction Kit, NEB).

### Plasmid challenge assay

ORF406 of conjugative plasmid pNOB8 was amplified using primers ORF406_fw/rv. Matching overhangs were introduced into pENTRY vectors downstream of a constitutive promoter (i.e truncated version of leader sequence of CRISPR array D in *Sa. solfataricus* P2) with primers pENTRY-ORF406_fw/rv. The resulting product was used as template in a touchdown PCR reaction (57°C, TΔ-0,2) for fusion with the ORF406 amplicon. ORF406 harbors a protospacer with a 5’NGA PAM exhibiting partial complementarity to spacer A53 of *Sa. solfataricus* P2^39^ which is absent from P1. Using primers OE-D2.109_fw/rv, this protospacer region was exchanged to match spacer 109 of CRISPR array D2 (D2.109) in P1 as described previously^24^. The insertion of an additional nucleotide directly downstream of the protospacer rendered the original PAM sequence dysfunctional impeding type I-A CRISPR-mediated interference. Additionally, we constructed a protospacer D2.109-a with an antitag (5’ CTTTCAAT) matching to the 5’ tag of the D2.109 crRNA using primer OE-D2.109-a_fw. The sequences of all primers used for construction of ORF406 target constructs are listed in Suppl. Tab. S6. Transformation of wildtype and deletions strain was carried out as described above, with the following alterations: prior to electroporation competent cells from both strains were adjusted to the same optical densities with 20 mM Sucrose, then three replicates of 50 μL cells mixed with 350 ng of plasmid DNA were set up for each construct. Cells without the addition of plasmid DNA were used as control. Following 1 hour recovery, each replicate was inoculated in 50 mL selective medium. After 41 hours (i.e. before cultures had started to grow), 45 μl of serial dilutions from 10^1^ to 10^-5^ from all cultures were spotted on selective gellan gum plates containing D(+)-Sucrose (0.2%, w/v) and NZ-Amine (0.1%, w/v).

### Adaptation experiment

M18 wildtype and KO2 were inoculated from glycerol stocks into prewarmed Brock medium S/T/U, 50 mL, pH 3.5, and incubated under shaking conditions at 77°C. As a preadaptation step, aliquots of both strains were then transferred to Brock medium G/T/U supplemented with D(+)-Glucose (Sigma Aldrich) instead of Sucrose at the same final concentration. A second aliquot was again transferred to Brock S/T/U. For the adaptation growth curve, 4 replicate cultures were set up for the following media compositions: Brock medium Sucrose (0.3%, w/v) with Uracil, Brock medium Glucose (0.3%, w/v) with Uracil, and standard complex Brock medium S/T/U. When cultures had reached late exponential growth, one replicate was transferred to 4 replicate flasks with fresh medium of the same composition until growth had stabilized for two consecutive transfers. At that point, 5 mL samples were taken at OD_600_=0.2 for RNA preparation. Throughout the experiment, cultures were inoculated to the same initial OD_600_.

### DNA preparation and genome mutation analysis

DNA was extracted as described in ref^24^. The sequencing was performed by the Next Generation Sequencing Facility at at Vienna BioCenter Core Facilities (VBCF), member of the Vienna BioCenter (VBC), Austria. Samples were sequenced (150 PE) sequencing on an Illumina MiSeq platform with a Westburg DNA preparation kit resulting in 300,000-500,000 reads per sample. Raw reads were downloaded from the VBCF website and checked using md5sum. FastQC^111^ version 0.11.8 and MultiQC^112^ version 1.8 were used to initially analyze sequences and the samtools package^113^ version 1.10 (using htslib 1.10^114^) along with bedtools ^115^ version 2.29.0 was used to convert file types. Trimmomatic^116^ version 0.36 trimmed adapter sequences and cropped reads using the options: SLIDINGWINDOW:5:20 LEADING:5 TRAILING:5 MINLEN:50 HEADCROP:12. The stand-alone program PrinSeq Lite^117^ version 0.20.4 was used to remove sequences < 38 bp and/or with quality scores < 30. Next, Bowtie2^118^ version 2.3.5.1 was used to align the remaining RNA sequences to the *Sa. solfataricus* strain P1 reference genome (NZ_LT549890.1) downloaded from the National Center for Biotechnology Information (NCBI). Mutations, insertions, and deletions were then found using breseq^119,120^ version 0.32.1 with default settings using the NCBI RefSeq annotation from 02/19/2020.

### RNA preparation and sequencing analysis

RNA was extracted from 5 mL of culture sampled at OD_600_=0.2 as described in ref^24^. Sample quality control, library preparation (150 PE) from rRNA-depleted samples (to approx. 50%) and sequencing on an Illumina NovaSeq platform was performed by Novogene (UK) Co., Ltd. Raw reads were downloaded from the sequencing facilities’ website and checked using md5sum. Adapter sequences were trimmed using Trimmomatic^116^ version 0.39 using the following settings: HEADCROP:12 SLIDINGWINDOW:4:15 LEADING:3 TRAILING:3 MINLEN:30. Reads were filtered for a minimum mean quality score of 30 using PrinSeq Lite^117^ version 0.20.4. Reads mapping to ribosomal RNA were removed from total reads with the sortmeRNA tool^121^ version 4.3.6 by supplying a curated reference file containing *Sa. solfataricus* P1 rRNA sequences. The remaining reads were aligned to the *Sa. solfataricus* P1 reference genome (NCBI: NZ_LT549890.1) using Hisat2^122^ version 2.2.1 with parameter --rna-strandness RF. The resulting sequence alignment files were converted to binary format using the samtools package^113^ version 1.20. featureCounts from the Subread package ^123^ version 2.0.6 assigned mapped reads to genomic features (feature and attribute type specified as “gene” and “ID”, respectively) using a General Feature Format (GFF) annotation file with RefSeq annotations downladed from the NCBI website on as a reference. To assess expression of transposase genes, a separate run with featureCounts was performed to inlcude reads mapping at multiple positions within the genome (‘-M’ option for multimapping reads). The output count matrix was used to perform differential expression analysis with the DESseq2 package^124^ version 1.42.1 in R^125^ version 4.3.3 with Rstudio^126^ version 2023.12.1+402. Default p-value adjustment in DESeq2 (Benjamini-Hochberg method) was used to determine relevant p-values (referred to as p-adj in result tables). Principal component analysis (PCA) was performed as described previoulsy^40^.

### Assignment of transporter families, arCOGs and transmembrane domains, and functional enrichment analysis

The BLAST tool integrated into the curated Transporter Classification Database (TCDB)^127^ was used to classify transporters. Archaeal Clusters of Orthologous Groups (arCOGs)^128^ were assigned to genes according to database version 2018. Genes without an arCOG assignment were annotated using eggNOG-mapper^129^ version 2.1.12. Genes encooding proteins with predicted transmembrane domains were identified using Phobius^130^. In both cases, a modified statistical hypergeometric test to determine functional categories significantly enriched for *Sa. solfataricus* (*P*-value < 0.05) among the up- or downregulated genes was conducted using the phyper function in R^40^.

