## Supplementary material for "The type III-B CRISPR-Cas System Affects Energy Metabolism and Adaptation in the Archaeon *Saccharolobus solfataricus*"

### Supplementary Figures and Tables

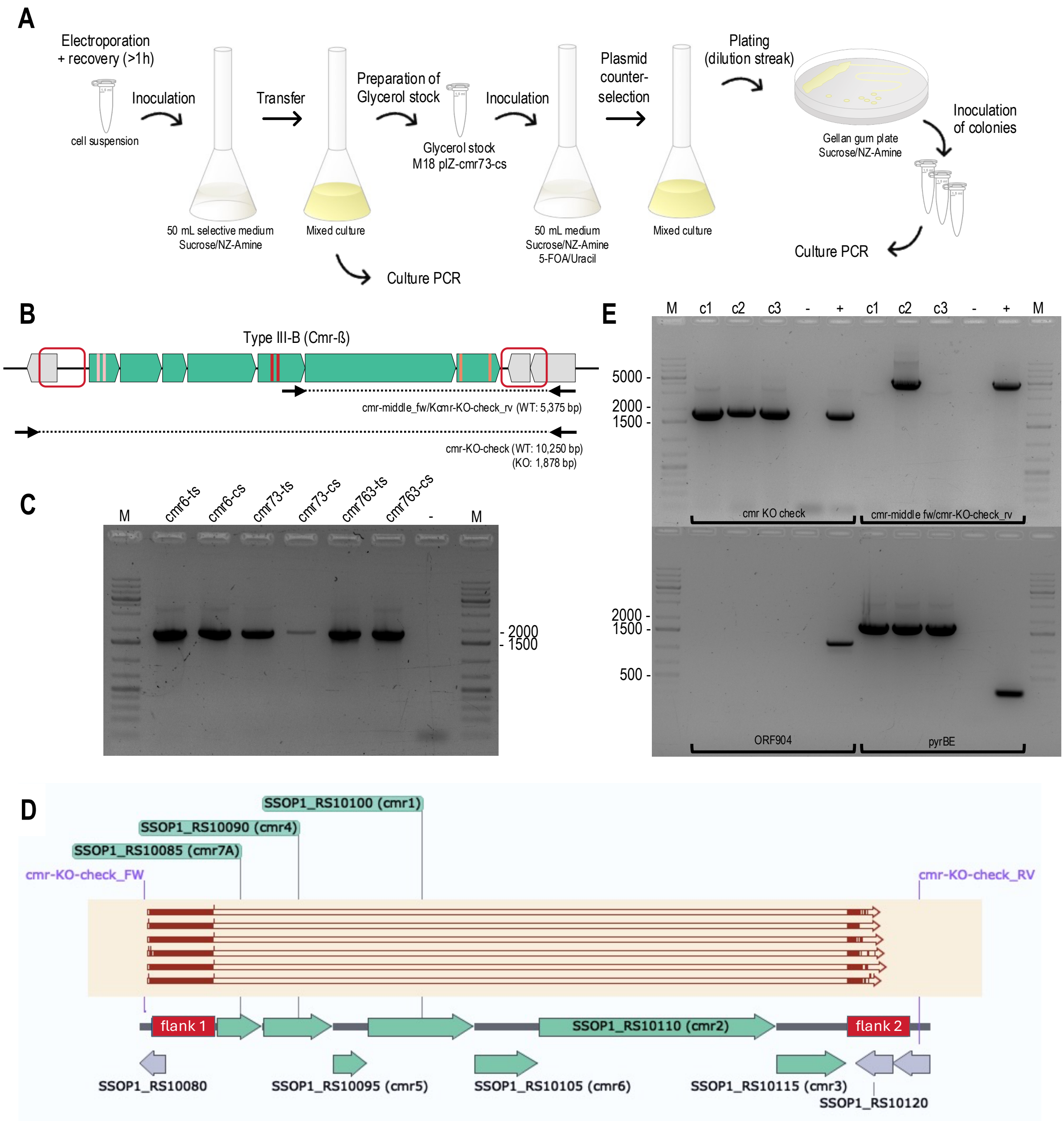

**Figure S1. Generation of *cmr-ß* knockout mutants in *Sa. solfataricus* P1. (A)** Workflow from electroporation to isolation of single mutant colonies. **(B)** Location of binding sites for primers used to verify *cmr-ß* knockout in culture PCR. **(C)** PCR products amplified with cmr-KO-check primers from mixed liquid cultures each transformed with one of six different plasmids indicated the desired knockout of the *cmr-ß* gene module. All plasmids harbored the same donor DNA sequences, but varying combinations of artifical spacers (one – three) targeting either coding or template strand (cs and ts, respectively) as indicated in (A). (-) – PCR negative control; M – 1 kb Plus DNA ladder (Thermo Scientific). **(D)** Sequencing of purified PCR product confirmed the scarless deletion for all transformants. Respective alignments to the reference genome are shown as red arrows (aligned - red shading, gap – no color) in the same order (top to bottom) as in (C). **(E)** For two colonies, c1 and c3 (subsequently renamed to KO1 and KO2, respectively), the sole presence of the *cmr-ß* mutant genotype was confirmed by PCR, while c2 exhibited a mixed genotype. Primer pairs ORF904_fw/rv (plasmid ORF904) and pyrBE_fw/rv were used to confirm the the successful counterselection against genome editing plasmids and the persistence of the IS element in the genomic *pyrEF* locus causing uracil auxothrophy. (-) – PCR negative control; (+) PCR positive control; M – 1 kb Plus DNA ladder (Thermo Scientific).

**Table S1.** Genes in the second deleted region in KO2.

| locus_tag | protein_ID | protein_product | arCOG_clu |
|---|---|---|---|
| SSOP1_RS17235 | WP_231918291.1 | hypothetical protein | N/A |
| SSOP1_RS17480 | WP_009992219.1 | hypothetical protein | N/A |
| SSOP1_RS14945 | WP_164497324.1 | ISH3 family transposase | arCOG03902 |
| SSOP1_RS14950 | WP_009993108.1 | sugar phosphate isomerase/epimerase family protein | arCOG01900 |
| SSOP1_RS14955 | WP_010923018.1 | ISH3-like element ISC1359 family transposase | arCOG03902 |
| SSOP1_RS17250 | WP_010924098.1 | hypothetical protein | N/A |
| SSOP1_RS17765 | WP_010924099.1 | hypothetical protein | N/A |
| SSOP1_RS14965 | WP_010924100.1 | transposase | arCOG09928 |
| SSOP1_RS16715 | WP_269454394.1 | hypothetical protein | N/A |
| SSOP1_RS17770 | WP_231918293.1 | fibronectin type III-like domain-containing protein | N/A |
| SSOP1_RS17775 | WP_231918295.1 | glycoside hydrolase family 3 C-terminal domain-containing protein | N/A |
| SSOP1_RS17780 | WP_231918343.1 | glycoside hydrolase family 3 N-terminal domain-containing protein | N/A |
| SSOP1_RS17785 | WP_231918297.1 | glycoside hydrolase family 3 N-terminal domain-containing protein | N/A |
| SSOP1_RS16720 | WP_049770880.1 | sugar phosphate isomerase/epimerase | arCOG01896 |
| SSOP1_RS14985 | WP_009990702.1 | dihydrodipicolinate synthase family protein | arCOG04172 |
| SSOP1_RS14990 | WP_009990700.1 | glycoside hydrolase family 2 TIM barrel-domain containing protein | arCOG07337 |
| SSOP1_RS17255 | WP_010924103.1 | DUF2139 domain-containing protein | N/A |
| SSOP1_RS15000 | WP_010924104.1 | DUF2139 domain-containing protein | N/A |
| SSOP1_RS15005 | WP_009990698.1 | GH116 family glycosyl hydrolase | arCOG03867 |
| SSOP1_RS15010 | WP_009990697.1 | SMP-30/gluconolactonase/LRE family protein | arCOG05370 |
| SSOP1_RS15015 | WP_009990696.1 | glucose 1-dehydrogenase | arCOG01459 |
| SSOP1_RS15020 | WP_010924107.1 | ABC transporter substrate-binding protein | arCOG08575 |
| SSOP1_RS15025 | WP_010924108.1 | ATP-binding cassette domain-containing protein | arCOG00184 |
| SSOP1_RS15030 | WP_010924109.1 | ABC transporter ATP-binding protein | arCOG00181 |
| SSOP1_RS15035 | WP_048054281.1 | ABC transporter permease | arCOG00748 |
| SSOP1_RS15040 | WP_010924111.1 | ABC transporter permease | arCOG00751 |
| SSOP1_RS15045 | WP_010924112.1 | Gfo/Idh/MocA family oxidoreductase | arCOG01622 |
| SSOP1_RS15050 | WP_009988433.1 | sugar phosphate isomerase/epimerase | arCOG01895 |
| SSOP1_RS15055 | WP_009988431.1 | alpha-glucosidase MalA | arCOG03663 |
| SSOP1_RS15060 | WP_009988429.1 | SGNH/GDSL hydrolase family protein | arCOG05121 |
| SSOP1_RS15065 | WP_063492843.1 | ABC transporter substrate-binding protein | arCOG01534 |
| SSOP1_RS15070 | WP_009988426.1 | aldo/keto reductase | arCOG01617 |
| SSOP1_RS15075 | WP_009988424.1 | ABC transporter ATP-binding protein | arCOG00181 |

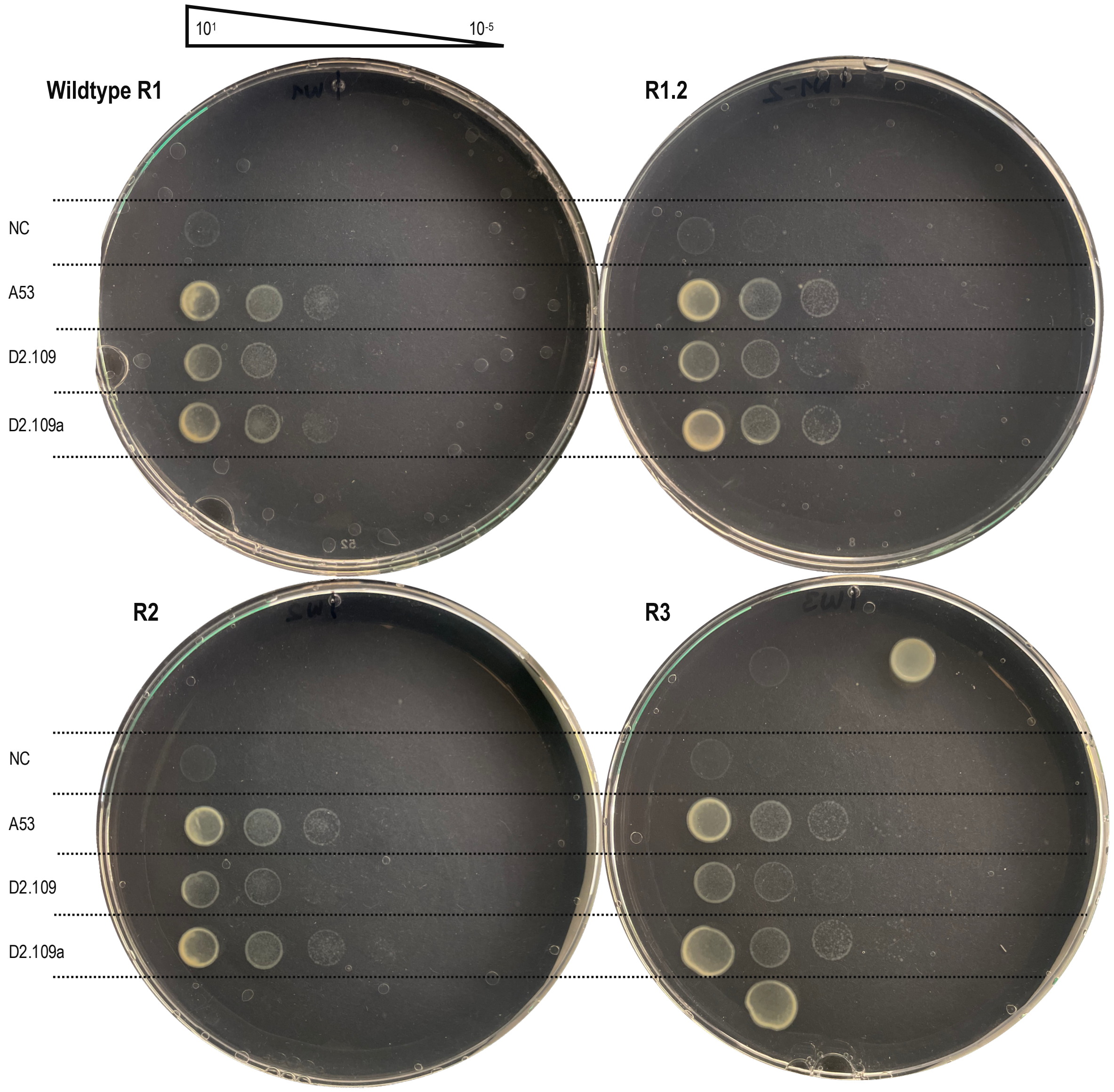


**Figure S2.** Plasmid challenge experiment with wildtype cells. Plates generated for all three biological replicates (R1-R3) during the spotting assay presented in main Figure 4 are shown. A second technical replicate plate was done for replicate 1 (R1.2).

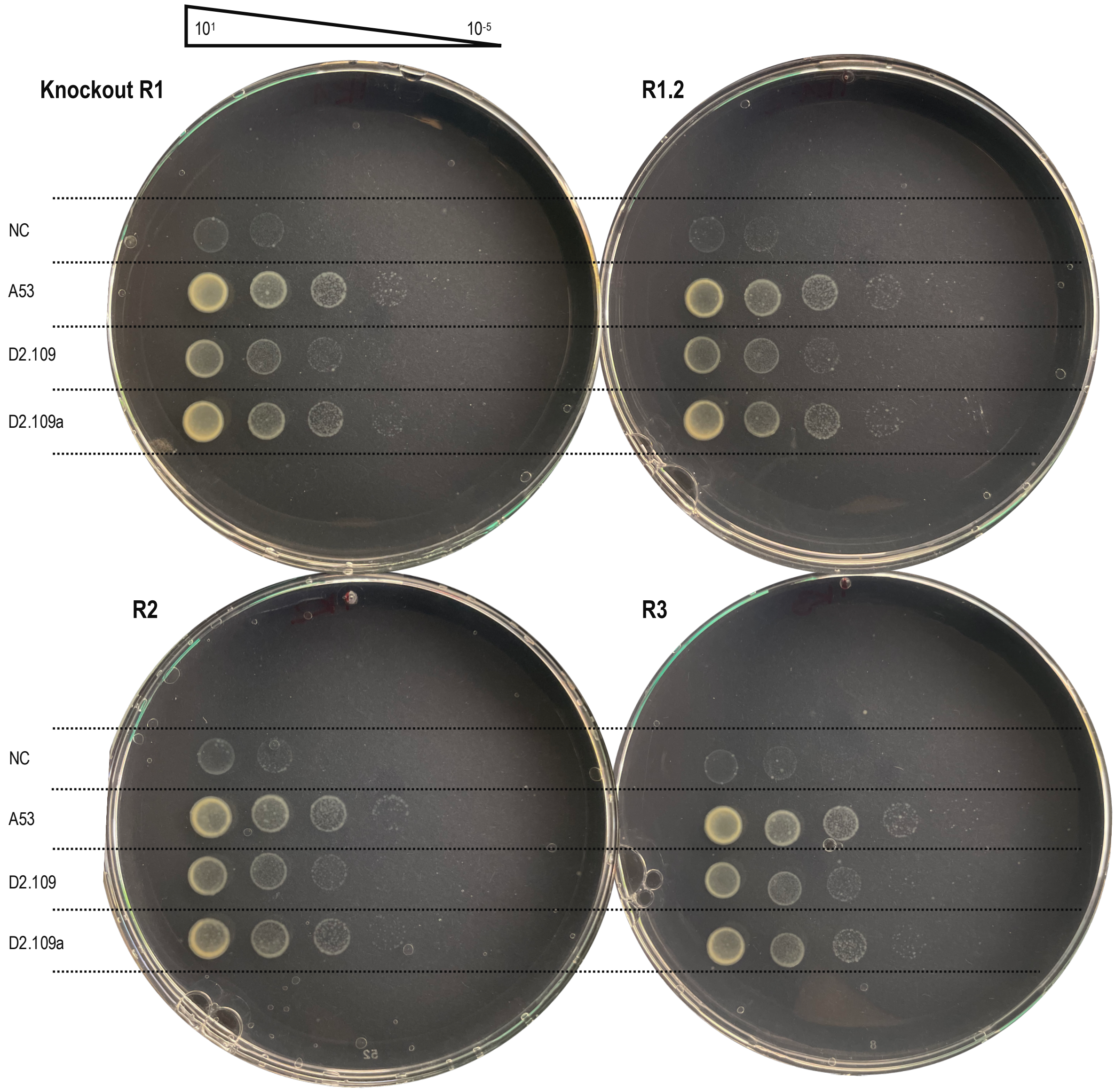


**Figure S3.** Plasmid challenge experiment with *cmr-ß* knockout mutant cells. Plates generated for all three biological replicates (R1-R3) during the spotting assay presented in main Figure 4 are shown. A second technical replicate plate was done for replicate 1 (R1.2).

**Table S2.** Top25 upregulated genes in *cmr-ß* knockout mutant strain KO1. Sorted based on log2-transformed fold change (FC) in gene expression when compared to wildtype. As a reference, log2-transformed TPM values are shown for both wildtype (WT TPM) and knockout (KO1 TPM). An adjusted *P* value (padj) < 0.001 was used to determine significance. Where available, transporter families or substrates are detailed in parenthesis in annotation column based on TCDB classification.

| locus_tag | WT TPM | KO1 TPM | FC | padj | protein_ID | gene | Annotation (arCOG database) |
|---|---|---|---|---|---|---|---|
| SSOP1_RS15825 | 8.17 | 11.52 | 3.56 | 0.00E+00 | WP_029552623.1 | - | Uncharacterized membrane protein |
| SSOP1_RS15675 | 6.63 | 8.82 | 2.41 | 8.32E-129 | WP_009991711.1 | - | MFS family permease<br>(2.A.1.9: The Phosphate: H+ Symporter (PHS) Family) |
| SSOP1_RS01655 | 6.54 | 8.44 | 2.11 | 1.78E-135 | WP_009990623.1 | nuoN | NADH dehydrogenase subunit N |
| SSOP1_RS15280 | 2.08 | 3.68 | 2.08 | 2.51E-26 | WP_063492868.1 | - | Uncharacterized membrane protein |
| SSOP1_RS15270 | 1.40 | 2.75 | 1.99 | 8.18E-10 | WP_063492866.1 | ccmB | ABC-type multidrug transport system, permease component |
| SSOP1_RS06715 | 4.98 | 6.69 | 1.94 | 8.30E-27 | WP_009990041.1 | - | Uncharacterized membrane protein |
| SSOP1_RS06710 | 5.77 | 7.44 | 1.89 | 3.80E-57 | WP_009990037.1 | appF | ABC-type oligopeptide transport system, ATPase component<br>(3.A.1.5.47: Peptide uptake porter) |
| SSOP1_RS01650 | 7.30 | 8.92 | 1.81 | 1.60E-71 | WP_009990622.1 | nuoL | NADH dehydrogenase subunit L |
| SSOP1_RS17770 | 4.16 | 5.69 | 1.79 | 9.64E-19 | WP_231918293.1 | - | - |
| SSOP1_RS15320 | 3.47 | 4.93 | 1.75 | 2.97E-27 | WP_063492874.1 | paaJ | Acetyl-CoA acetyltransferase |
| SSOP1_RS15215 | 4.47 | 5.98 | 1.75 | 3.82E-26 | WP_063492857.1 | - | DsrE/DsrF/DrsH-like family peroxiredoxin |
| SSOP1_RS01645 | 5.62 | 7.12 | 1.70 | 1.05E-26 | WP_009990621.1 | nuoK | NADH dehydrogenase subunit 4L (K,kappa) |
| SSOP1_RS06995 | 3.43 | 4.75 | 1.66 | 7.90E-08 | WP_010923318.1 | thiC | Thiamine biosynthesis protein ThiC |
| SSOP1_RS06700 | 6.70 | 8.13 | 1.64 | 1.07E-59 | WP_009990034.1 | dppC | ABC-type dipeptide/oligopeptide/nickel transport system, permease component<br>(3.A.1.5.47: Peptide uptake porter) |
| SSOP1_RS06705 | 6.09 | 7.50 | 1.63 | 9.48E-39 | WP_009990035.1 | dppD | ABC-type dipeptide/oligopeptide/nickel transport system, ATPase component<br>(3.A.1.5.47: Peptide uptake porter) |
| SSOP1_RS15390 | 5.01 | 6.40 | 1.62 | 8.39E-45 | WP_063492885.1 | kdaD | 2-keto-3-deoxyarabinoate/xylonate dehydratase |
| SSOP1_RS15025 | 3.33 | 4.60 | 1.57 | 4.89E-16 | WP_010924108.1 | appF | ABC-type oligopeptide transport system, ATPase component<br>(3.A.1.5.8: Maltose MalEFGK) |
| SSOP1_RS15395 | 1.94 | 3.06 | 1.55 | 1.75E-07 | WP_063492886.1 | - | MFS family permease<br>(2.A.1.15: The Aromatic Acid:H+ Symporter (AAHS) Family) |
| SSOP1_RS01640 | 6.93 | 8.27 | 1.55 | 9.04E-52 | WP_009990620.1 | nuoJ | NADH dehydrogenase subunit J |
| SSOP1_RS01635 | 7.58 | 8.91 | 1.55 | 2.92E-51 | WP_009990619.1 | nuoI | NADH dehydrogenase subunit I |
| SSOP1_RS03205 | 6.31 | 7.64 | 1.54 | 4.04E-63 | WP_009991168.1 | purD | Phosphoribosylamine-glycine ligase |
| SSOP1_RS03210 | 6.12 | 7.45 | 1.54 | 1.80E-45 | WP_009991170.1 | purM | Phosphoribosylaminoimidazole (AIR) synthetase |
| SSOP1_RS15345 | 3.53 | 4.79 | 1.54 | 7.35E-19 | WP_063492878.1 | tatC | Sec-independent protein secretion pathway component TatC |
| SSOP1_RS15305 | 3.17 | 4.38 | 1.51 | 1.30E-15 | WP_063492871.1 | - | Uncharacterized protein |
| SSOP1_RS01630 | 7.55 | 8.84 | 1.49 | 1.69E-36 | WP_009990618.1 | nuoH | NADH dehydrogenase subunit H |

**Table S3.** Top25 upregulated genes in *cmr-ß* knockout mutant strain KO2. Sorted based on log2-transformed fold change (FC) in gene expression when compared to wildtype. As a reference, log2-transformed TPM values are shown for both wildtype (WT TPM) and knockout (KO2 TPM). An adjusted *P* value (padj) < 0.001 was used to determine significance. Where available, transporter families or substrates are detailed in parenthesis in annotation column based on TCDB classification.

| locus tag | WT TPM | KO2 TPM | FC | padj | protein ID | gene | annotation (arCOG database) |
|---|---|---|---|---|---|---|---|
| SSOP1_RS14475 | 0.76 | 2.54 | 2.81 | 1.75E-10 | WP_010924134.1 | - | IS4 transposase |
| SSOP1_RS06715 | 4.98 | 7.41 | 2.50 | 5.30E-50 | WP_009990041.1 | - | Uncharacterized membrane protein |
| SSOP1_RS06710 | 5.77 | 8.12 | 2.40 | 2.16E-143 | WP_009990037.1 | appF | ABC-type oligopeptide transport system, ATPase component (3.A.1.5.47: Peptide uptake porter) |
| SSOP1_RS15280 | 2.08 | 4.08 | 2.38 | 3.27E-23 | WP_063492868.1 | - | Uncharacterized membrane protein |
| SSOP1_RS06705 | 6.09 | 8.33 | 2.29 | 8.63E-152 | WP_009990035.1 | dppD | ABC-type dipeptide/oligopeptide/nickel transport system, ATPase component |
| SSOP1_RS01655 | 6.54 | 8.64 | 2.15 | 2.24E-137 | WP_009990623.1 | nuoN | NADH dehydrogenase subunit N |
| SSOP1_RS15130 | 2.45 | 4.40 | 2.13 | 1.67E-17 | WP_063492845.1 | araT | ABC-type sugar transport system, permease component (3.A.1.1.14-AraS: arabinose, fructose, xylose) |
| SSOP1_RS06700 | 6.70 | 8.68 | 2.02 | 1.62E-115 | WP_009990034.1 | nppC | ABC-type dipeptide/oligopeptide/nickel transport system, permease component (3.A.1.5.47: Peptide uptake porter) |
| SSOP1_RS15320 | 3.47 | 5.23 | 1.90 | 1.22E-36 | WP_063492874.1 | paaJ | Acetyl-CoA acetyltransferase |
| SSOP1_RS01650 | 7.30 | 9.13 | 1.87 | 7.09E-85 | WP_009990622.1 | nuoL | NADH dehydrogenase subunit L |
| SSOP1_RS15345 | 3.53 | 5.27 | 1.86 | 5.74E-34 | WP_063492878.1 | tatC | Sec-independent protein secretion pathway component TatC |
| SSOP1_RS15135 | 2.63 | 4.23 | 1.81 | 6.49E-15 | WP_063492846.1 | araU | ABC-type sugar transport system, permease component (3.A.1.1.14-AraS: arabinose, fructose, xylose) |
| SSOP1_RS05370 | 3.67 | 5.33 | 1.79 | 2.20E-04 | - | - | - |
| SSOP1_RS01645 | 5.62 | 7.37 | 1.79 | 2.27E-35 | WP_009990621.1 | nuoK | NADH dehydrogenase subunit 4L (K,kappa) |
| SSOP1_RS03210 | 6.12 | 7.86 | 1.79 | 8.66E-81 | WP_009991170.1 | purM | Phosphoribosylaminoimidazole (AIR) synthetase |
| SSOP1_RS15270 | 1.40 | 2.67 | 1.72 | 2.96E-07 | WP_063492866.1 | ccmB | ABC-type multidrug transport system, permease component |
| SSOP1_RS15390 | 5.01 | 6.67 | 1.72 | 1.52E-65 | WP_063492885.1 | kdaD | 2-keto-3-deoxyarabinoate/xylonate dehydratase |
| SSOP1_RS10720 | 3.21 | 4.78 | 1.71 | 2.89E-09 | WP_009990146.1 | marR | Transcriptional regulator, MarR family |
| SSOP1_RS06695 | 7.61 | 9.22 | 1.66 | 3.71E-84 | WP_063492780.1 | dppB | ABC-type dipeptide/oligopeptide/nickel transport system, permease component (3.A.1.5.47: Peptide uptake porter) |
| SSOP1_RS10725 | 3.70 | 5.24 | 1.66 | 1.42E-17 | WP_009990147.1 | - | Membrane component of uncharacterized ABC transporter |
| SSOP1_RS03205 | 6.31 | 7.92 | 1.65 | 4.76E-90 | WP_009991168.1 | purD | Phosphoribosylamine-glycine ligase |
| SSOP1_RS15360 | 7.06 | 8.67 | 1.64 | 5.30E-95 | WP_063492880.1 | ypwA | Zn-dependent carboxypeptidase, M32 family |
| SSOP1_RS01640 | 6.93 | 8.51 | 1.63 | 1.15E-82 | WP_009990620.1 | nuoJ | NADH dehydrogenase subunit J |
| SSOP1_RS15305 | 3.17 | 4.64 | 1.62 | 1.18E-18 | WP_063492871.1 | - | Uncharacterized protein |
| SSOP1_RS15300 | 6.20 | 7.71 | 1.55 | 1.90E-105 | WP_063492870.1 | - | Extracellular solute-binding protein with Ig-fold domain |

**Table S4.** List of 20 genes that were significantly upregulated in *cmr-ß* knockout mutant KO2 across all three conditions in the adaptation experiment (Sucrose-Tryptone, Sucrose, Glucose) in comparison to wildtype samples.

| locus tag | protein ID | gene | protein product | arCOG cluster | annotation (arCOG database) | func. category |
|---|---|---|---|---|---|---|
| SSOP1_RS00905 | WP_009990418.1 | hemC | hydroxymethylbilane synthase | arCOG04299 | Porphobilinogen deaminase | H |
| SSOP1_RS02750 | WP_009991040.1 | hmgB | hydroxymethylglutaryl-CoA synthase | arCOG01767 | 3-hydroxy-3-methylglutaryl CoA synthase | I |
| SSOP1_RS02755 | WP_009991041.1 | aact | thiolase family protein | arCOG01278 | Acetyl-CoA acetyltransferase | I |
| SSOP1_RS02760 | WP_009991042.1 | DUF35 | Zn-ribbon domain-containing OB-fold protein | arCOG01285 | OB-fold domain and Zn-ribbon containing protein, possible acyl-CoA-binding protein | R |
| SSOP1_RS02765 | WP_010923027.1 | hmgA | hydroxymethylglutaryl-CoA reductase (NADPH) | arCOG04260 | Hydroxymethylglutaryl-CoA reductase | I |
| SSOP1_RS06140 | WP_063492735.1 | cutA | aerobic carbon-monoxide dehydrogenase large subunit | arCOG01167 | Aerobic-type carbon monoxide dehydrogenase, large subunit CoxL/CutL homolog | C |
| SSOP1_RS06150 | WP_009992007.1 | cutB | xanthine dehydrogenase family protein subunit M | arCOG01926 | Aerobic-type carbon monoxide dehydrogenase, middle subunit CoxM/CutM homolog | C |
| SSOP1_RS06345 | WP_231918216.1 | - | nickel-dependent hydrogenase large subunit | - | - | - |
| SSOP1_RS07390 | WP_009988400.1 | cas3a2 | CRISPR-associated endonuclease Cas3'' | arCOG01443 | HD superfamily nuclease | V |
| SSOP1_RS07550 | WP_010923402.1 | cas6 | CRISPR-associated endoribonuclease Cas6 | arCOG01439 | CRISPR-Cas system related protein, RAMP superfamily Cas6 group | V |
| SSOP1_RS09060 | WP_269454399.1 | - | MFS transporter | arCOG00130 | MFS family permease | G |
| SSOP1_RS10570 | WP_009989795.1 | - | protease pro-enzyme activation domain-containing protein | arCOG03669 | Subtilase family protease | O |
| SSOP1_RS14685 | WP_010924079.1 | soxI | hypothetical protein | arCOG06033 | Predicted subunit of heme/copper-type cytochrome/quinol oxidase | C |
| SSOP1_RS14690 | WP_010924080.1 | soxH | cytochrome c oxidase subunit II | arCOG01235 | Heme/copper-type cytochrome/quinol oxidase, subunit 2 | C |
| SSOP1_RS14695 | WP_173645069.1 | soxG | cytochrome bc complex cytochrome b subunit | - | - | - |
| SSOP1_RS14700 | WP_009992566.1 | soxF | Rieske 2Fe-2S domain-containing protein | arCOG01720 | Rieske Fe-S protein | C |
| SSOP1_RS14705 | WP_009992568.1 | soxE | sulfocyanin | arCOG03700 | Sulfocyanin | C |
| SSOP1_RS14710 | WP_173645070.1 | soxM | cbb3-type cytochrome c oxidase subunit I | arCOG01237 | Heme/copper-type cytochrome/quinol oxidase, subunit 1 and 3 | C |
| SSOP1_RS15280 | WP_063492868.1 | - | hypothetical protein | arCOG08496 | Uncharacterized membrane protein | S |
| SSOP1_RS17650 | WP_014511669.1 | - | hypothetical protein | - | - | - |

**Table S5.** Top25 genes contributing to variance explained by principal component 2 (PC2) in direction of *cmr-ß* knockout samples (most negative) when compared to wildtype samples across all three conditions of adaptation experiment.

| locus tag | protein_ID | gene | protein product | arCOG cluster | annotation (arCOG database) | func. category |
|---|---|---|---|---|---|---|
| SSOP1_RS10600 | WP_009989783.1 | treY | malto-oligosyltrehalose synthase | arCOG02955 | Maltooligosyl trehalose synthase | G |
| SSOP1_RS12565 | WP_009989901.1 | - | Fis family transcriptional regulator | arCOG01842 | Sterol carrier protein | I |
| SSOP1_RS13020 | WP_009989316.1 | - | hypothetical protein | arCOG08529 | Predicted permease (3.A.1.128: The SkfA Peptide Exporter (SkfA-E) Family) | R |
| SSOP1_RS00330 | WP_009988870.1 | mho1 | AmmeMemoRadiSam system protein B | arCOG01728 | Predicted class III extradiol dioxygenase, MEMO1 family | R |
| SSOP1_RS01720 | WP_009990638.1 | nusG | transcription elongation factor Spt5 | arCOG01920 | Transcription termination/antitermination protein NusG | K |
| SSOP1_RS12760 | WP_010923916.1 | - | hypothetical protein | arCOG07325 | Uncharacterized membrane protein | S |
| SSOP1_RS00325 | WP_009988869.1 | - | hypothetical protein | arCOG05890 | Mevalonate kinase related protein | R |
| SSOP1_RS04190 | WP_048054188.1 | - | hypothetical protein | arCOG07186 | PIN domain | V |
| SSOP1_RS00925 | WP_009990424.1 | hcaD | (2Fe-2S)-binding protein | arCOG01299 | NAD(FAD)-dependent dehydrogenase | R |
| SSOP1_RS01655 | WP_009990623.1 | nuoN | NADH-quinone oxidoreductase subunit NuoN | arCOG01540 | NADH dehydrogenase subunit N | C |
| SSOP1_RS00900 | WP_009990416.1 | hemL | glutamate-1-semialdehyde 2,1-aminomutase | arCOG00918 | Glutamate-1-semialdehyde aminotransferase | H |
| SSOP1_RS01650 | WP_009990622.1 | nuoL | proton-conducting transporter membrane subunit | arCOG01539 | NADH dehydrogenase subunit L | C |
| SSOP1_RS00905 | WP_009990418.1 | hemC | hydroxymethylbilane synthase | arCOG04299 | Porphobilinogen deaminase | H |
| SSOP1_RS07230 | WP_010923362.1 | - | hypothetical protein | arCOG02487 | Cell surface protein, a component of a putative secretion system | M |
| SSOP1_RS01565 | WP_009990605.1 | dinG | ATP-dependent DNA helicase | arCOG00770 | Rad3-related DNA helicase | K |
| SSOP1_RS05605 | WP_009989736.1 | - | MFS transporter | arCOG00130 | MFS family permease (2.A.1.14: The Anion:Cation Symporter (ACS) Family) | G |
| SSOP1_RS12560 | WP_009989902.1 | acrR | TetR/AcrR family transcriptional regulator | arCOG02647 | Transcriptional regulator, TetR/AcrR family | K |
| SSOP1_RS02765 | WP_010923027.1 | hmg1 | hydroxymethylglutaryl-CoA reductase (NADPH) | arCOG04260 | Hydroxymethylglutaryl-CoA reductase | I |
| SSOP1_RS06715 | WP_009990041.1 | - | hypothetical protein | arCOG08395 | Uncharacterized membrane protein | S |
| SSOP1_RS01645 | WP_009990621.1 | nuoK | NADH-quinone oxidoreductase subunit K | arCOG03073 | NADH dehydrogenase subunit 4L (K,kappa) | C |
| SSOP1_RS09310 | WP_009992714.1 | - | nucleotidyltransferase domain-containing protein | arCOG01204 | Minimal nucleotidyltransferase | V |
| SSOP1_RS13555 | WP_009988643.1 | ugpE | ABC transporter permease subunit | arCOG00158 | ABC-type sugar transport system, permease component (3.A.1.1: The Carbohydrate Uptake Transporter-1 (CUT1) Family) | G |
| SSOP1_RS00565 | WP_014511489.1 | upsB | archaellin/type IV pilin N-terminal domain-containing protein | arCOG07276 | Pilin/Flagellin, FlaG/FlaF family | N |
| SSOP1_RS07550 | WP_010923402.1 | cas6 | CRISPR-associated endoribonuclease Cas6 | arCOG01439 | CRISPR-Cas system related protein, RAMP superfamily Cas6 group | V |
| SSOP1_RS09585 | WP_009992018.1 | - | DUF1286 domain-containing protein | arCOG05320 | Predicted membrane-bound metal-dependent hydrolase | R |

**Table S6.** Construction PCR primers used in this study. Recognition sequences of restriction enzymes are underlined.

| Construction Primer | Sequence (5’ to 3’) |
|---|---|
| OE-cmr7-cs_fw | ATTTGACTGACATGTTGTCCATTCTGATTGATAATCTCTTATAGAATTGA |
| OE-cmr7-cs_rv | TGGACAACATGTCAGTCAAATCGATGATACTTTCAATTCTATAAGAGATT |
| OE-cmr6-cs_fw | CCTCGTCCTAGTAGTTATTTCAAAGTCTAGATAATCTCTTATAGAATTGA |
| OE-cmr6-cs_rv | GAAATAACTACTAGGACGAGGGCTTTAATCTTTCAATTCTATAAGAGATT |
| OE-cmr3-cs_fw | CTTATCCACCTTACCCATTGTAACATAATGATAATCTCTTATAGAATTGA |
| OE-cmr3-cs_rv | ACAATGGGTAAGGTGGATAAGATAAGCTTCTTTCAATTCTATAAGAGATT |
| OE-cmr7-ts_fw | GTACCTACTTTTGATCCTTGTGATTACGTGATAATCTCTTATAGAATTGA |
| OE-cmr7-bstr_rv | ACAAGGATCAAAAGTAGGTACTAGTTCGCCTTTCAATTCTATAAGAGATT |
| OE-cmr6-bstr_fw | GGGAAATAGCTGAACTTGTCTCAAGGAACGATAATCTCTTATAGAATTGA |
| OE-cmr6-bstr_rv | AGACAAGTTCAGCTATTTCCCTCTTAAATCTTTCAATTCTATAAGAGATT |
| OE-cmr3-bstr_fw | GTTAATGTTTAGAAGTCAGGGGGAATTTGGATAATCTCTTATAGAATTGA |
| OE-cmr3-bstr_rv | CCCTGACTTCTAAACATTAACGGTTCTACTTTCAATTCTATAAGAGATT |
| MOE_fw | AGAATTATCGCCCAGAACAAATTTCTGATAATCTCTTATAGAATTGAAAG |
| MOE_rv | GTTAGTTCACCCACCGACAAATACAACTTTCAATTCTATAAGAGATTATC |
| flank1_fw_SalI | AAGCGTCGACCCAATAACCAACAATTAGAGTCCC |
| flank1_rv | CATAATACAAAAGTTCTTACGCGCTT |
| flank2_fw | AATGCTATTACATTCCTACGTTTGTCCC |
| flank2_rv_EagI | GAAACGGCCGAATGCGCTATTCCGACTCACAG |
| ORF406_fw | GACTTATCATCAACCCAGTAATG |
| ORF406_rv | ATGCCCAGCATAGGATTTTGTTTTC |
| pENTRY-ORF406_fw | AAACAAAATCCTATGCTGGGCATTGAGGGTTTATTAATCTCTTACTATTT |
| pENTRY-ORF406_rv | CATTACTGGGTTGATGATAAGTCAGCCTGCTTTTTTGTACAA |
| OE-ORF406_fw | GTAGTAATGGAGGAGATAGACGTTAAACAGTTGGTGGAAAAGTCCGAAT |
| OE-ORF406_rv | GCCGTACTTCTCAAGCTGGTACTTCAAGATGCTTTTCAGCTC |
| OE-D2.109_fw | TTCTGGTCCTTATACGAAATCGAGCTGAAAAGCATC |
| OE-D2.109-a_fw | TTCTGGTCCTTATACGAAACTTTCAATGAGCTGAAAAGCATC |
| OE-D2.109_fw | TTCTGGTCCTTATACGAAATCGAGCTGAAAAGCATC |
| OE-D2.109_rv | GTATAAGGACCAGAACGGCAATACCCAAACTGTTGATTCGGACTTTTCCA |

| Primer | Sequence (5’ to 3’) |
|---|---|
| cmr-KO-check_fw | TGCGTTTATGTATGATTGTGCG |
| cmr-KO-check_rv | GAGCTGATTTTACTGCCCGC |
| cmr-middle_fw | CTAGGCGAGCAAGATGCTGAGG |
| pyrB_fw | TCATCTCTGGTCAAGTCAAGC |
| pyrE_rv | GAATGCCGATAAGGGAACTC |
| ORF904_fw | ACAAGAAGAACGGGGGTG |
| ORF904_rv | ACCTCTTCAGCAATCGCCT |

Supplementary Notes

Notes on the additional deleted region in KO2:

This region contains 33 genes mainly encoding transposases and several genes related to sugar metabolism (Suppl. tab. 1). The later include multiple glycoside hydrolases, the substrate-binding subunit as well as ATPase component of a maltose ABC-transporter, and another ABC-transporter of the PepT family (3.A.1.5.8: Maltose MalEFGK)[1].

Responses of central metabolic pathways in the knockout grown on complex media:

Genes involved in sulfur metabolism (sulfite:acceptor oxidoreductase, SAOR; thiosulfate:quinone oxidoreductase, TQO; Heterodisulfide reductase, Hrd), transport and trafficking (*dsrE, tusA*) were mostly significantly downregulated in the type III deletion mutants, in accordance with the fact that autotrophic growth on elemental sulfur, sulfite, thiosulfate or tetrathionate compounds has never been shown for this species and therefore any putative contribution to the quinone pool is not significant[2]. A slight difference between both KO mutants can be observed in the non-significant slight upregulation of TQO and DoxBCE in KO1, which may indicate an attempt to convert thiosulfate oxidation to proton export.

Two different uptake systems for phosphate were upregulated in response to the deletion, namely the PhoT ABC transporter (3.A.1.7) as well as another high affinity transporter of the phosphate: H+ Symporter Family (2.A.1.9). This observation would be in agreement with the general hypothesis of essential nutrient uptake and membrane-related processes being upregulated in the *cmr-ß* knockout mutants.

Upregulation of a regulatory network of genes implicated in fatty acid metabolism was observed upon type III-B deletion. In contrast to bacteria and eukaryotes, fatty acids are not present in membrane lipids of archaea[3]. Although many archaeal genomes harbor genes related to fatty acid metabolism, their exact role and directionality of the pathway in archaea remain enigmatic[4,5]. In *S. acidocaldarius*, FadRSa was shown to directly regulate its own fatty acid metabolism gene cluster negatively. Furthermore, transcriptomic analysis of *fadR* deletion mutants also hinted at an inversely correlated, indirect effect on sulfur metabolism and cytochrome-containing membrane complexes[6]. Another study extended the experimentally investigated FadR regulon to include several more genes using a machine learning approach[7]. Excitingly, this analysis predicted the inclusion of the CRISPR-Cas associated transcriptional regulator Csa3 (saci_1992) into the FadR regulatory network. Csa3 and type III-D system appeared to be negatively correlated with the expression of the FadR gene cluster in response to organic solvent stress[8]. In our datasets, we could not detect a significant change in expression levels for any of the three transcriptional regulators associated with CRISPR loci in *Sa. solfataricus* P1.

Notes on the interpretation of the lipid biosynthesis transcriptional patterns:
The membrane composition in archaea has been shown to change dynamically as a homeoviscous adaptation to various environmental stressors such as temperature, pH, and nutrient availability, in order to achieve a balance between stability/rigidity and permeability at any given condition[9]. Therefore, when attempting to integrate our observations on the transcriptional profiles of the lipid biosynthesis pathway genes in a general interpretation framework, we also have to take into account possible connections to regulatory networks of sensing environmental stress, as the cell membrane is one of the primary cellular compartments to exhibit changes in response to adverse environmental conditions.

The structural features of GDGTs in archaeal membranes (e.g. number of cyclopentyl rings, reduction state and type of head groups) have been extensively studied in response to various stressors such as temperature, pH and pressure[9–12]. Recent studies have also revealed an underlying relationship between the membrane composition and the energetic status of the cell, as defined by the rate of metabolic activity, growth stage and availability of electron donors/acceptors[12–16]. The modulation of environmental parameters such as temperature and pH in various studies have nevertheless produced lipid patterns not always interpretable based on the level of energy stress experienced by the cell[12].

Ring Index (RI) is negatively correlated with growth temperature in *Sulfolobales*, where deviations from the optimal growth temperature have been shown to result in either increased or decreased RIs under higher and lower temperatures respectively[12,13,15], consistent with a homeoviscous adaptation. Responses to pH deviations have been shown to be more complex, involving both ring synthesis, modification of head groups and type of backbone[10]. While the RI generally decreased in either increased or decreased pH with respect to optimum conditions[12,17], a closer look revealed an increase in GDGTs with ≥ 4 rings at pH 2 and an increase in GDGTs with 3 and 4 rings at pH 4[12], with presumably opposite effects on membrane functioning, concomitant with the differential regulation of the two GDGT ring synthases in response to temperature and pH stress[18]. Calditol-linked GDGTs have also been shown to be essential for growth of *Sulfolobus acidocaldarius* during acid stress[19], due in part to the increased hydrogen bonding capacity of calditol vs glycerol, thus leading to a further reduction in ion permeability[10,19].

Similar changes to the GDGT profiles in model Thermoproteota (Sulfolobales and Nitrososphaeria) and Euryarchaeota were observed as a result of a decreased energy supply, either in the form of nutrient depletion during late growth stages or a controlled limitation in the energy supply[12–16,20,21]. Both these conditions resulted in an increase in the RI, presumed to lead to a decrease in membrane ion permeability, and thus reduction in cellular energy needs. An additional underlying mechanism proposed is that ring formation prevents double-bond reduction of the isoprenoid chains by GGR, a reaction that consumes electrons, thereby conserving reducing power under conditions where energy flow is not optimal[13].

Observations on the differential expression of transposase genes:
In *Sa. solfataricus*, a transposon-derived ncRNA was shown to influence transcription levels of a phosphate transporter gene through antisense regulation in trans (high affinity phosphate transporter)[22]. Interestingly, the overexpression of a ncRNA stemming from an IS200/605 element caused an increased-growth phenotype in *Halobacterium salinarum*[23].

Transcriptional responses during the adaptation of the knockout from complex to minimal media:
Although generally less genes were significantly differentially expressed in the standard sucrose-tryptone condition in the deletion mutant compared to the initial transcriptome, the vast majority of upregulated genes (73 %) was consistent with the previous dataset (Fig. 9A). Most importantly, this included the NADH dehydrogenase (complex I) and terminal oxidase SoxEFGHIM (complex III), the arabinose ABC transporter as well as other transporters mentioned above, type III-D CRISPR-Cas genes, genes involved in lipid biosynthesis and thiamine transport and biosynthesis.

In the combined PCA of all conditions (Fig. 10B), PC1 was separating the complex from the defined media conditions thereby explaining 49 % of the observed variance. Interestingly, PC2 accounting for approximately 12 % of the explained variance seemed to separate the wildtype from the *cmr-ß* knockout samples.

When looking at the top25 genes that drive the separation along PC2 (Tab. S5) (i.e., genes that show higher expression in *cmr-ß* knockout compared to wildtype samples), we found genes encoding subunits of complex I and proteins partaking in tetrapyrrole biosynthesis (SSOP1_RS00900 - *hemL*, SSOP1_RS00905 - *hemC*), as well as various transporters (CUT1 family ABC transporter, etc). In addition, genes encoding proteins involved in lipid biosynthesis (SSOP1_RS02765 - *hmgA*, SSOP1_RS00325 - Mevalonate kinase related protein), two uncharacterized mebrane proteins (SSOP1_RS12760 14 TMs, next to APC family permease, SSOP1_RS06715 located downstream of peptide uptake porter of PepT family) and a component of a putative secretion system (SSOP1_RS07230) are seemingly major drivers of variance along PC2 according to our statistical analysis.

Interestingly, the list also included two neighboring genes with convergent orientation, one of which encodes a putative sterol-binding domain protein (SSOP1_RS12565, arCOG01842, COG3255). The second encodes for a transcriptional regulator of the TetR/AcrR family (SSOP1_RS12560), which are one-component signal transduction systems consisting of an N-terminal HTH DNA-binding and a C-terminal ligand binding domain. Characterized members of this family have been shown to interact with several different ligands and regulate a variety of physiological processes, such as antibiotic resistance, cell signalling, fatty acid metabolism etc[24]. The homolog present in *S. acidocaldarius*, termed FadRSa, has been chracterized recently and was shown to regulate fatty acid metabolism genes[6]. It acts as a repressor of a 30kb gene cluster which includes lipases and β-oxidation enzymes, however the operational direction is not understood. FadRSa was shown to specifcally bind acyl-CoA molecules and consequently dissociate from bound DNA. Binding sites were bioinformatically predicted for a similar gene cluster in *Sa. solfataricus* P2, which also included the *fadR* promoter itself[6]. In our datasets, we found the corresponding genes in *Sa. solfataricus* P1 to be mostly upregulated except for in the glucose condition.
